# Newfoundland and Labrador: A mosaic founder population of an Irish and British diaspora from 300 years ago

**DOI:** 10.1101/2022.04.01.486593

**Authors:** Edmund Gilbert, Heather Zurel, Margaret E. MacMillan, Sedat Demiriz, Sadra Mirhendi, Michael Merrigan, Seamus O’Reilly, Anne M. Molloy, Lawrence C. Brody, Walter Bodmer, Richard A. Leach, Roderick E. M. Scott, Gerald Mugford, Ranjit Randhawa, J. Claiborne Stephens, Alison L. Symington, Gianpiero L. Cavalleri, Michael S. Phillips

## Abstract

The founder population of Newfoundland and Labrador (NL) is a unique genetic resource, in part due to geographic and cultural isolation, where historical records describe a migration of European settlers primarily from Ireland and England to NL in the 18th and 19th centuries. Whilst its historical isolation, and increase prevalence of certain monogenic disorders, have been appreciated, the fine-scale genetic structure and ancestry of the population has not been well described. Understanding the genetic background on which functional, disease causing, genetic variation resides on would aid informed genetic mapping efforts in the Province. Here, we leverage dense genome-wide SNP data on 1,807 NL individuals to reveal fine-scale genetic structure in NL that is clustered around coastal communities and correlated with Christian denomination. We show that the majority of NL European ancestry can be traced back to the south-east and south-west of Ireland and England, respectively. We date a substantial population size bottleneck approximately 10-15 generations ago in NL, associated with increased haplotype sharing and autozygosity. Our results elucidate novel insights into the population history of NL and demonstrate evidence of a population conducive to further genetic studies and biomarker discovery.

**Significance Statement:** Newfoundland and Labrador (NL) has been identified as a founder population, though evidence of its magnitude and subsequent isolation is unclear. Here, analysis of 1,807 NL individuals demonstrates population structure associated with geographical isolation in coastal communities and religious denomination (Catholic or Protestant Christian). Further, NL European ancestry primarily descends from settlers from south-east Ireland and south-west England. This history is associated with increased sharing of longer haplotypes in NL, and NL-specific drift in some communities more than others, providing strong evidence of a founder event occurring about 10-15 generations ago. This study elucidates the detailed population structure of NL and shows enrichment for otherwise low frequency functional variants due to genetic drift useful for potential future biomarker discovery studies.

## Introduction

Newfoundland and Labrador (NL) is the most eastern Canadian province. It is comprised of Labrador on the Canadian mainland, and the island of Newfoundland located in the north Atlantic (Figure 1). The predominantly European-ancestry population has expanded to approximately 520,000 (Statistics Canada) today from the initial 25,000 European settlers who came to NL in the 18th and 19th centuries. These European settlers who migrated to NL were predominantly of Irish and English origin1. Historical records suggest that these European migrants came from Irish, predominantly Catholic, communities in the southwestern Irish counties of Waterford, Wexford, south Kilkenny, southeast Tipperary, and southeast Cork2. In the case of the English, the mainly Protestant settlers can be traced back to the counties of Dorset and Devon as well as the fishing ports such as Dartmouth, Plymouth, or Southampton in southwestern England2,3. Prior to and after European settlement of the region several Indigenous Peoples including the maritime Archaic peoples, Mi’kmaq, the Innu and the Inuit, and Beothuk, inhabited areas within the modern Province.4,5. The modern population of NL includes peoples of Indigenous ancestry, primarily the Innuit, Innu, and Mi’kmaq. The Innuit and Innu are mainly located within Labrador, and the Mi’kmaq are found within Newfoundland, both with some level of admixture with people of European ancestry3.

**Figure 1.**
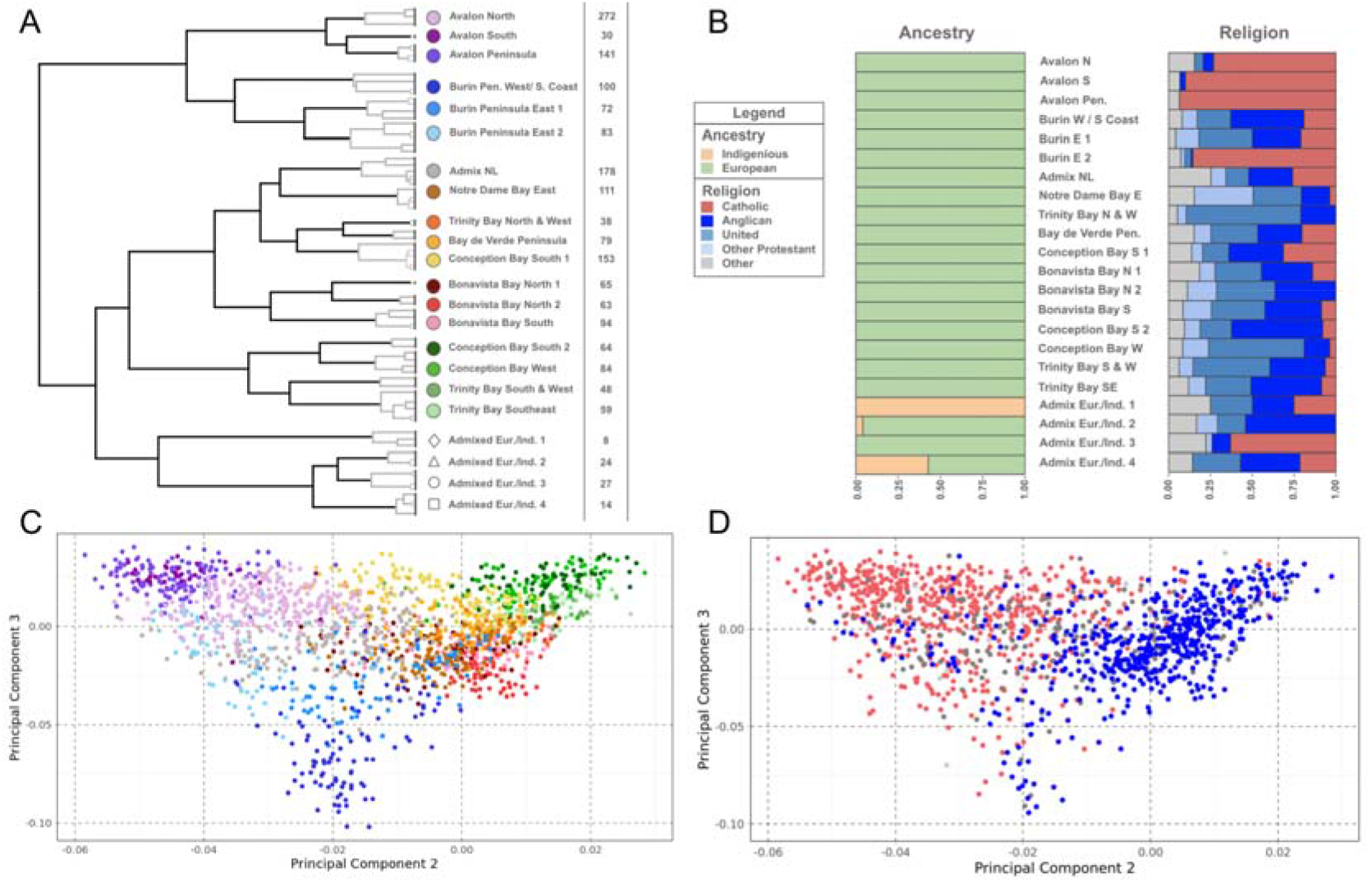
Genetic structure of Newfoundland and Labrador. (A) Dendrogram of the *fineSTRUCTURE* final MAP state, showing 22 summarising clusters of NL membership. Solid branches indicate the 22 cluster branches, with grey branch-lines showing merged clusters within each of the 22 clusters. *fineSTRUCTURE* clusters are colour and shaped coded to reflect grouping on adjacent branches, with labels reflecting common geographic birthplace of members’ grandparents. Cluster sizes shown to the right of cluster labels. (B) The proportions of genetic ancestry groups and religious background in each of the 22 clusters, shown in the order of A. (C) The second versus the third principal components of the *ChromoPainter* co-ancestry matrix, with individual colour and shape coded according to *fineSTRUCTURE* cluster membership. (D) The second versus the third principal components of the *ChromoPainter* co-ancestry matrix, with individual points colour coded to religious background; red indicating Christian Catholic, blue indicating Christian Protestant, and grey indicating other or unknown. All panels were plotted using the statistical computing language R^58^ and the packages ggplot2 and rworldxtra.

Migration of Catholic Irish and Protestant English settlers to NL peaked in the mid to late 18th century3. These migrants who settled into coastal communities (known as outports) were socially and geographically isolated from one another, rarely intermarrying and so experienced subsequent “private” bottlenecks. This cultural and geographical isolation is mirrored in the genetic landscape of NL. Previous studies using limited genetic data, found evidence of genetic isolation and elevated inbreeding coefficients4,6,7 as well as conflicting evidence of extended linkage disequilibrium1,3,8, features that have been successfully leveraged to identify both recessive and dominant Mendelian traits9-13. Recently analysis of genome-wide SNP-array genotype data from NL individuals3 confirmed evidence of genetic isolation in the NL population, and that the broad NL population structure could be described in terms of; i) indigenous American ancestry and, ii) Catholic versus Protestant background.

NL is therefore potentially of great value to genetic mapping efforts, similar to that realised from other genetically isolated populations14,15. Isolated populations with a history of population bottlenecks are valuable communities for genetic mapping efforts as genetic drift increases the frequencies of rare clinically relevant variants as well as increasing average haplotype length and general homogeneity15, which are characteristics of founder events. As isolated populations are increasingly leveraged in the study of rare or ultra-rare genetic variation16, an appreciation of their fine-scale genetic landscape is needed to account for the increased stratification of rarer genetic variation17,18. Although haplotype-based methods19 have demonstrated fine-scale genetic structure in many populations including the ancestral source populations of Britain and Ireland20-23, such approaches have not been applied within the context of NL. Furthermore, applying haplotype-based approaches to the NL context could also reveal insights into the population’s demographic history and isolation24-27.

Given this context, we aimed to explore the non-Indigenous (European) settlement of the present NL population in unprecedented detail, studying a large sample of individuals with NL ancestry, together with ancestry source references from Britain and Ireland. Leveraging data from 1,807 individuals with European founder ancestry from the Newfoundland and Labrador Genome Project (NLGP) we set out to; i) characterise the fine-scale population structure in NL using haplotype-based methods and investigate how this structure relates to ancestry, religion, and geography, ii) quantify the proportions of British and Irish ancestry in NL and map these to their regional sources in Britain and Ireland using ancestry references21,28,29, and finally iii) characterise the extent that a history of bottlenecks has had on the haplotype diversity of NL compared to ancestral sources in Britain and Ireland.

## Results

### Newfoundland Genetic Structure

To establish the fine-scale genetic structure and origins of the European settler NL population and post individual-genotype and SNP-genotype quality control (QC) (see Methods), we had access to data on 2,446 participants from the Newfoundland and Labrador and Genome Project (NLGP – Sequence Bio). To sample the genetic landscape of NL before modern economic migration in the latter 20th century onwards, we used principal component analysis (PCA). We projected the NLGP individuals onto world-wide genetic variation to identify NLGP individuals who occupied the same ancestry space as either European or Indigenous American ancestry references from HGDP or 1KGP3 (see Figures S1.1-1.2). Having identified a sample of “NL ancestry”, we further performed sample and marker QC (see Methods), leaving a core dataset of 1,807 NL individuals (the “NL1,807” dataset) and 685,221 common SNPs for further analysis.

To investigate the fine-scale genetic structure of NL, we performed haplotype-based clustering using *fineSTRUCTURE* to cluster individuals based on their haplotype sharing, as quantified by the *ChromoPainter* co-ancestry matrix. *fineSTRUCTURE* analysis identified 22 discrete clusters which summarise *fineSTRUCTURE*’s 74 clusters from its final maximum a posteriori (MAP) state. This k=22 level of clustering combines smaller, difficult to interpret, clusters together to summarise the predominant fine-scale structure present in NL (Figure 1A). The 22 clusters are further grouped in a dendrogram, organising clusters that share excess haplotypes together on shared branches. Most clusters show geographical stratification (Figure 2 and Figures S2.1-2.6 for individual plots) as well as religious. The clusters exhibit substantial genetic differentiation, as measured by FST (average FST: 0.00206, min FST: 0. 00016, max FST: 0.00429; Table S1). This mean differentiation is an order of magnitude higher than what is found between equivalent *fineSTRUCTURE*-cluster estimates from Ireland or England (FST: 0.0003, or 0.0003 respectively)21.

**Figure 2.**
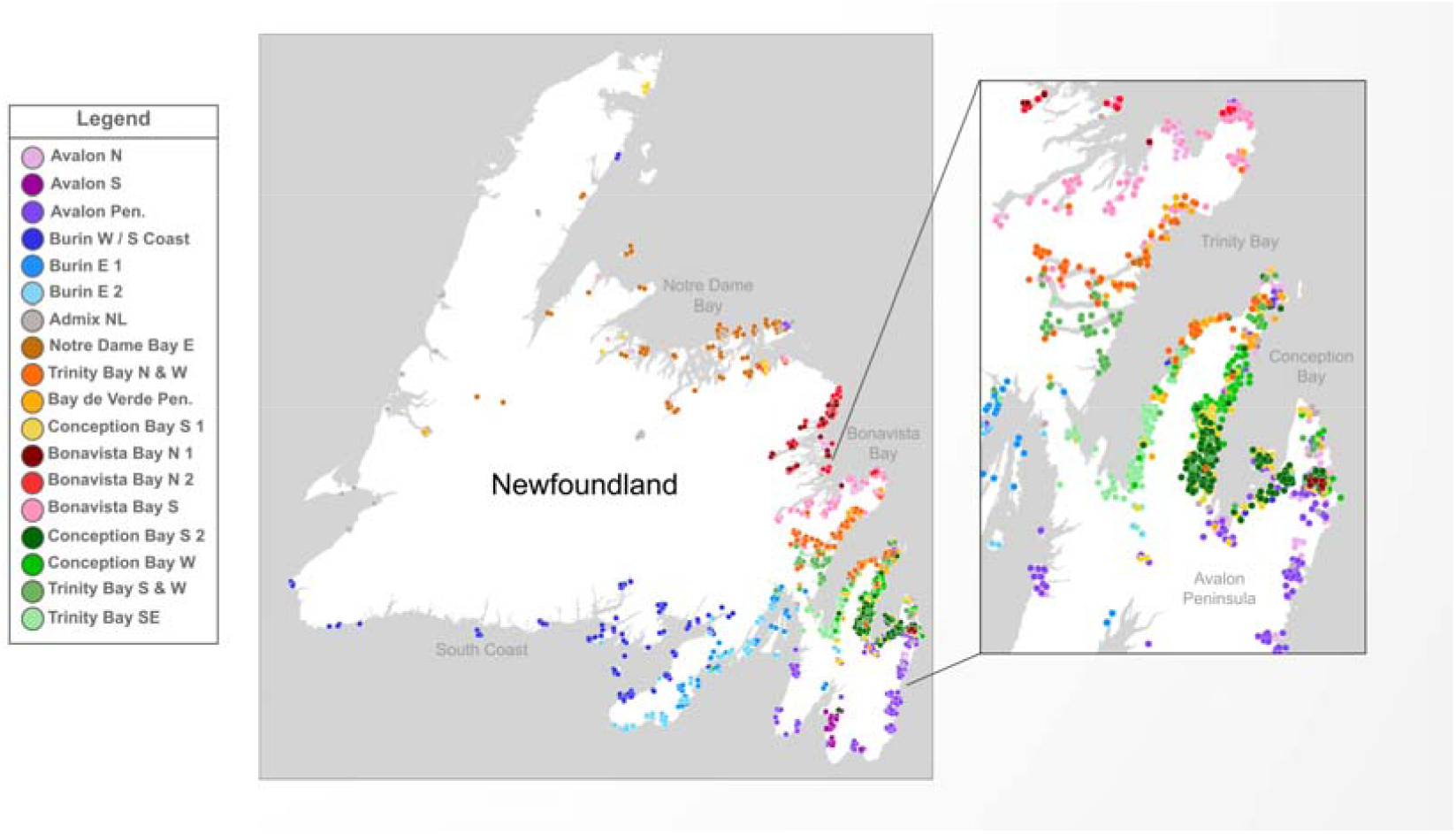
Genetic landscape of Newfoundland and Labrador. Map of the grandparental birthplaces of individuals with colour and shape coded according to fineSTRUCTURE cluster. A small jitter has been introduced to aid legibility and preserve anonymity. An insert shows individual details of the Trinity and Conception Bays. Panel was plotted within photoshop, with geography boundary data sourced from Tableau.

The first split in the *fineSTRUCTURE* dendrogram separates individuals in the south-east of NL with grandparents predominantly from either Burin or the Avalon peninsula into six clusters (Figure 2). Indeed, individuals with ancestry from south-eastern NL appear to be genetically distinct from the rest of NL in PCA as well as clustering (Figure 1C). When we compared the proportion of individuals in each *fineSTRUCTURE* cluster associating with various Christian denominations, we found four out of the six clusters (Figure 1B, 1D) show high proportions of Catholic background. There are significant differences of religious background between genetic NL clusters as measured by Χ2 test (p = 0.00049), in agreement with previous work3. These differences are largely driven by Catholic membership within the Avalon peninsula (the Avalon Pen. and Avalon N clusters), whose residuals contribute approximately 22% of the overall Χ2 statistic. Furthermore, PCA of the *ChromoPainter* coancestry matrix supported this observation by separating individuals with a Catholic or Protestant background along PC2, echoing previous observations3 (Figure 1D, Figure S2.6).

The other major branch in the *fineSTRUCTURE* dendrogram separated out clusters of the remaining NL individuals and those with putative Indigenous ancestry (Figure 1A & 1B). These individuals cluster with world-wide Indigenous American ancestry references in PCA (Figure S1.1), separate from other NL individuals along the first PC in NL-only PCA (Figure S2.7), and present higher proportions of American ancestry components in a supervised ADMIXTURE30 analysis using 1KGP3 ancestry references (Figure S2.8). As the focus of this study was to characterise the genetic structure of NL arising from the non-indigenous European settlers and given the lack of appropriate North American Indigenous references in addition to the small number of participants in these clusters (4% of the cohort), we made no further effort to characterise indigenous ancestry in subsequent analyses, focussing analysis on the remaining 18 clusters.

Beyond the south-eastern branch of the dendrogram, clusters demonstrate fine-scale structure among predominantly Protestant communities. These clusters align strikingly with the geographic features of individual NL bays. For example, the north-eastern Trinity and Conception Bays exhibit population structure not only between one another (Figure 2) but also within the same bay (Figures S2.1-2.6). This picture is similar in the northern bays of Bonavista and Notre Dame as well. FST distances between these clusters show substantial differentiation between the neighbouring communities, consistent with genetic isolation (Table S1). While most of these clusters are largely Protestant in religious background, there are subtle differences in denomination (Figure 1B). The large cluster of individuals with grandparents from Trinity Bay, Trinity Bay N&W, show a high proportion (68%) of United Church of Canada background, as does the Conception Bay W cluster (57%), compared to an average of 24% elsewhere. Considering the United Church of Canada post-dates settlement this could reflect specific communities preferentially favouring one denomination.

Elsewhere, the Admix NL cluster contains individuals with recent genealogies from across the island, and whose average copying vectors from the *ChromoPainter* coancestry matrix suggests haplotype sharing with different clusters across NL (Figure S2.9). Further, using the dimension-reduction methods UMAP31 and t-SNE32, these individuals co-locate across the space of NL individuals (Figure S2.10-11). We infer, therefore, that the Admix NL cluster likely captures individuals with a mix of ancestors from across NL grouped together through *fineSTRUCTURE*’s clustering algorithm. This mixed cluster could be due to modern economic movement in the 20th century, where communities prior to the 20th century were typically isolated. We observe a similar copying profile in the Avalon N cluster (Figure S2.9), which could represent the urban admixture in the metropolitan area of St John’s, the province’s largest city which is located to the north of the Avalon peninsula.

### Newfoundland Settler Ancestry

The relative ancestry contributions from Irish and British source populations to different NL communities is largely unknown, but assumed to correlate with historical records of settlement2 and Catholic/Protestant religious background3,33. To elucidate this, we used IBD-segment sharing patterns (see Methods) on a combined dataset of 1,807 individuals of NL ancestry and 4,408 ancestry reference individuals from Britain and Ireland, 1,808 of whom have geographic annotation. We first identified sub-communities within Ireland and Britain using IBD sharing network clustering34 (see Methods). We identified 26 IBD-clusters across Ireland and Britain that confirm previous *fineSTRUCTURE*-based clustering patterns (Figure 3A). With this set of regional Irish and British reference clusters, we then leveraged an extension of a previously reported nnls-based approach20 to model the proportions of IBD sharing between “target” and “source” clusters as estimated ancestry profiles.

**Figure 3.**
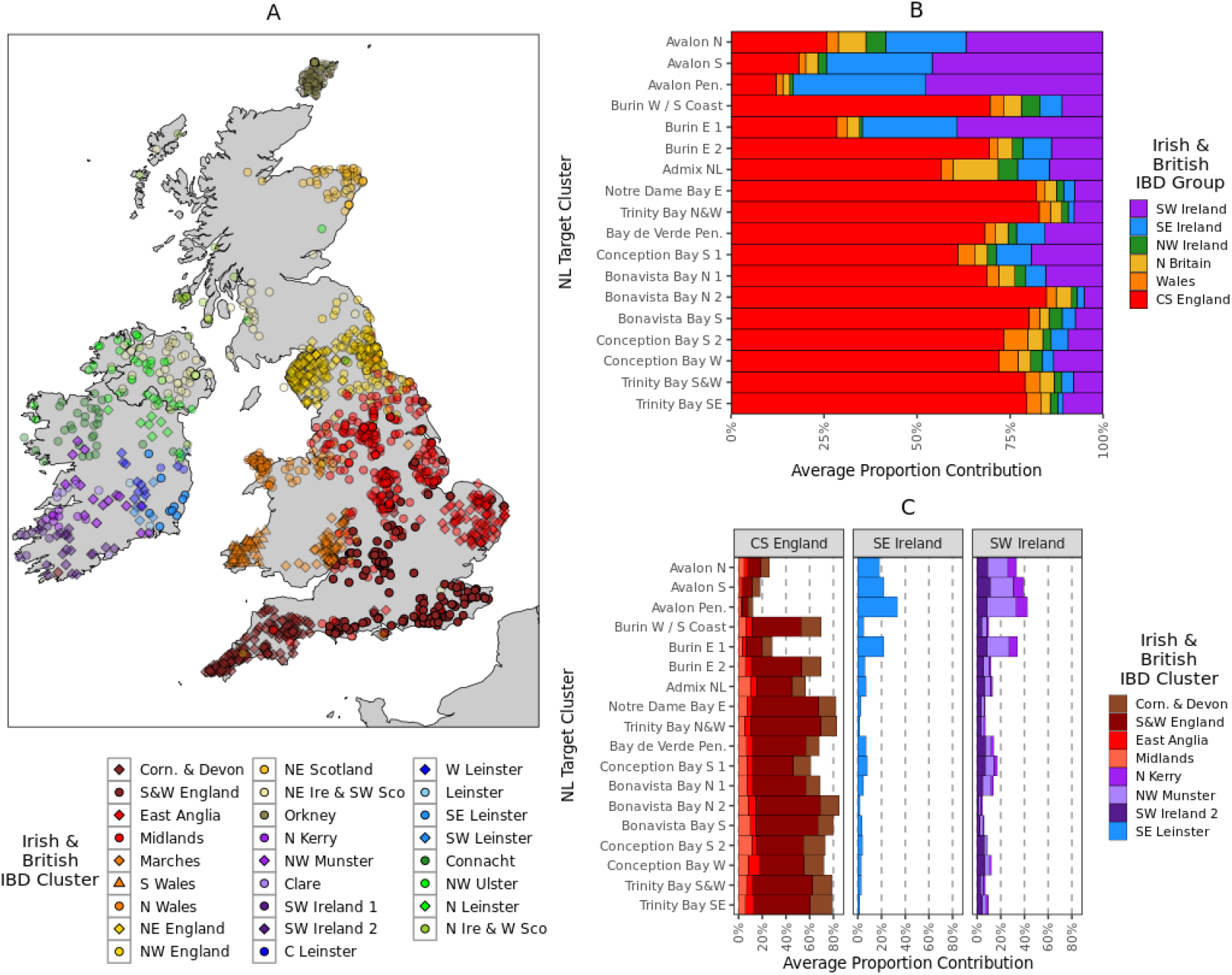
Irish and British ancestry in Newfoundland and Labrador. (A) Map of Irish^21^ and British^20^ individuals placed according to recent mean ancestral birthplace, colour and shape coded according to IBD-network clustering. Colours are used to group hierarchically related clusters together (see Methods). A slight transparency for individual points has been introduced proportional to the maximal proportion that each cluster contribute to any one NL cluster, with great contributors having less transparency. (B) The summed proportions of the estimated ancestry proportions calculated from Irish or British IBD-segments shared with NL. Contributions from individual IBD clusters are summed together in groups according to Irish or British region. (C) Individual Irish or British IBD cluster contributions to NL ancestry proportions, showing only clusters that contribute >5% to any one NL cluster. All panels were plotted using the statistical computing language R^58^ and the packages ggplot2 and rworldxtra.

Historical records suggest that the European settlers of NL were predominantly from south-eastern Ireland and south-western England2. To formally test this mixture, we only considered IBD-segment sharing between NL individuals and Irish or British reference individuals, further considering IBD segments >3 or < 15 cM in length (thereby capturing recent genealogical relationships). Our rationale being that if different NL clusters carry different proportions of Irish or British ancestry, this will differentiate in the amount of IBD segments that they carry – thus creating different “copying profiles”.

We estimated the average IBD contribution from each Irish or British source IBD-cluster to each NL *fineSTRUCTURE*-cluster target, as estimated by the nnls method. We first summed the contributions from general Irish or British regions, such as the south-west of England, or southern Ireland (Figure 3B). This showed substantially different profiles within NL, which are largely driven by either English or southern Irish ancestry contributions. The NL clusters with a high proportion of Catholic background (typically from the Avalon peninsula) are significantly associated with increasing Irish contribution (Welch Two-Sample test p= 0.009), and in general a high proportion of individuals with a Catholic background are associated with a high proportion of southern Irish ancestry (pearson r2=0.95, p=7e-12) (Figures S4.1 and S4.2).

Next, we investigated if any specific IBD-cluster of Irish or British individuals were driving these English/Irish signals. Considering individual source clusters that contribute substantially to any one NL target cluster (i.e., > 5%), we indeed find specific regional affinity within the English and Irish ancestry components (Figure 3C). The English component is driven by the S&W England cluster, as well as Corn. & Devon. The Irish component is largely driven by clusters with individuals who have recent genealogical ancestry (i.e., from the 1850s) from the Wexford and Waterford regions of southern Ireland, primarily N&W Munster and SE Leinster. Both these English and Irish contributions to Irish-British IBD-segment sharing with NL are strikingly supportive of historical records which show the migrants who migrated to NL can be traced back to communities from these regions in Britain and Ireland2. Moreover, the contributions from each individual Irish/British cluster to the Irish or British ancestry profile in NL are in similar proportions across NL clusters. This is suggestive of a single source of the Irish and British ancestry in NL, i.e., that the Irish or British ancestry in NL is not from multiple waves from different regions in Ireland or Britain.

This sharing signal in the southwest of NL of Irish haplotypes from the southeast of Ireland is further supported in an “unsupervised” form of the nnls method where we consider each NL or Irish-British cluster as a mixture of shared IBD from any other NL or Irish-British cluster (Figure S4.3). This analysis shows that whilst most NL clusters predominantly share IBD-contributions from other NL clusters, reflective of their shared ancestry, some clusters such as Burin E 1 or Avalon Pen. still present substantial ancestry contributions from the N&W Munster and SE Leinster clusters. Furthermore, we tested this mixture of Irish and British haplotypes with the fastGLOBETROTTER algorithm, whose mixture model agrees with an Irish ancestry source best represented by N&W Munster and SE Leinster (see Supplemental Data 5).

### Evidence of Population Bottleneck and Homogeneity

Existing literature from historical2, clinical1,9-13, and population genetic studies3 suggests evidence of a population bottleneck in the European settlement of NL. We set out to characterise the magnitude of this founder effect in NL using the NL1,807 dataset, comparing the NL profile to European source populations. We additionally explored the evidence of enrichment of rare or uncommon alleles in NL compared to Ireland or Britain.

We sought to estimate the historical effective population size (Ne) of NL to provide insight to past population bottlenecks, which would be expected to leave evidence of a reduction in Ne. We utilised the detected IBD-segments in our combined NL, Ireland, and Britain dataset and applied IBDNe25 to IBD-segment sharing within NL clusters >100 individuals in size. For comparison, we estimated using the same methods, Ne in the six broad regions of Ireland and Britain (Figure 4A-B). Both Ireland and Britain, and NL have experienced a period of exponential growth within the last 10 generations, consistent with previous estimates of other European ancestry populations35,36. Within Ireland and Britain, in general England has a higher Ne than Wales, Scotland, or Ireland, and Orkney. British estimates are consistent with previous estimations from the PoBI dataset36. Within NL we detect a consistent reduction of ancestral population size 15-10 generations ago across tested clusters. This population size reduction is several orders of magnitude lower to Irish or British equivalents at a comparable time-period. Ne estimates prior to this reduction are much larger than Irish or British estimates and have wide intervals, perhaps due to the mixed Irish-British ancestry in NL, or the impact of the settlement-bottleneck masking previous demographic profiles. NL regions associated with inter-regional admixture such as Avalon N or Admix NL have higher Ne estimates within the past 10 generations which would support evidence of an intra-NL admixture history.

**Figure 4.**
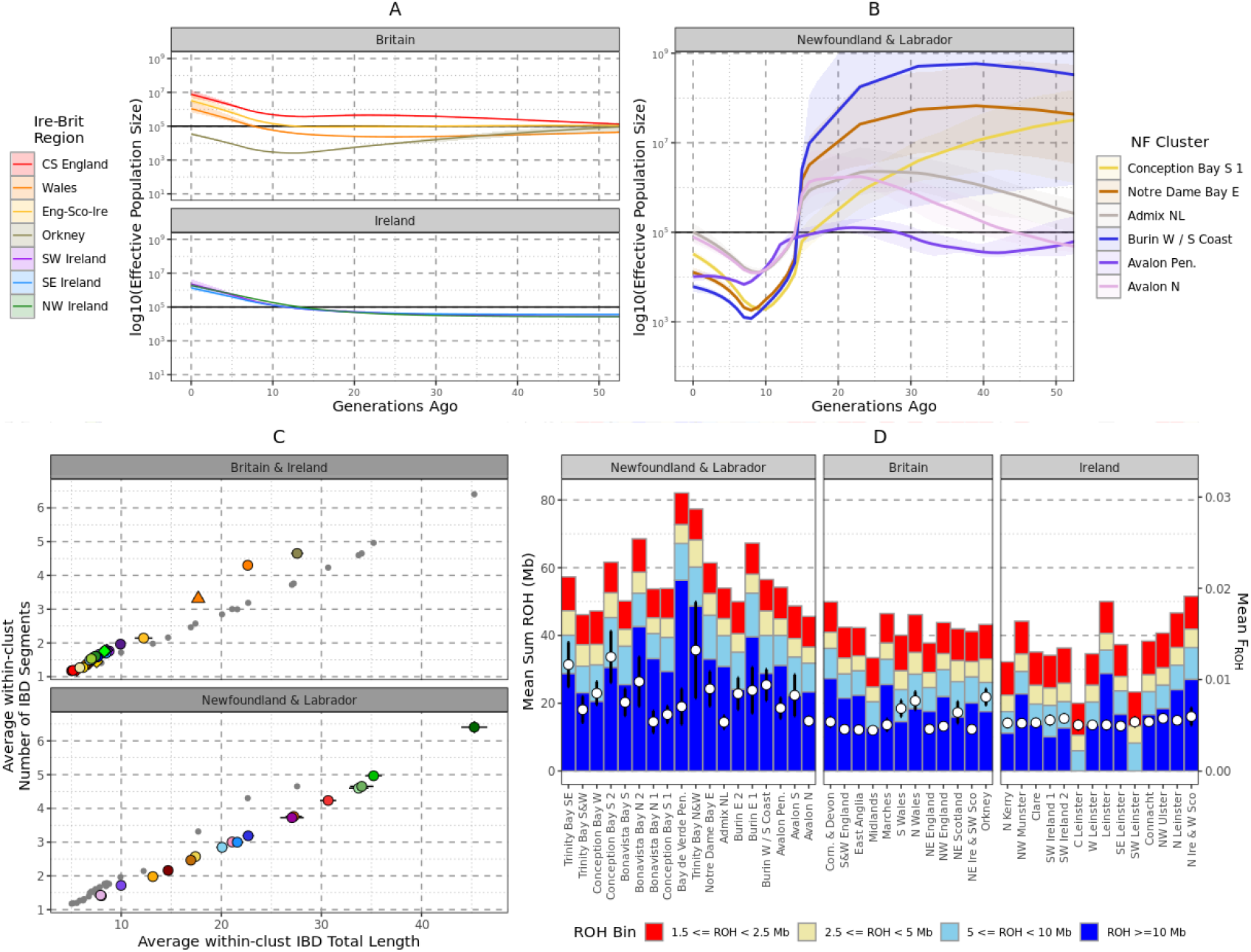
Evidence of bottleneck with Newfoundland and Labrador. (A) The estimated historical Ne for Irish and British regions using IBD-segments and the IBDNe tool. Shading shows the 95% confidence intervals. (B) The estimated historical Ne for NL clusters > 100 individuals in membership size. Shading indicates 95% confidence intervals. (C) The individual mean total length of Identity-by-Descent (IBD) segments shared with another individual placed in the same cluster, versus the mean number of IBD segments shared with individuals placed in the same cluster. Clusters are colour and shaped coded in the same system as D. Irish and British clusters are plotted separately from NL clusters for legibility, and grey points indicate non-Irish and British (or non-NL) clusters. Error bars show the 95% confidence intervals. (D) The average total length of ROH within each NL or Irish and British cluster. The mean total length of ROH > 1.5 Mb for each cluster is shown by a hollow white circle (left y axis), or the proportion of the genome in ROH > 1.5Mb in length (FROH – right y axis). Also shown are average total lengths of ROH in four length bins: 1) 1.5 Mb ≤ ROH ≤ 2.5 Mb, 2) 2.5 Mb ≤ ROH ≤ 5 Mb, 3) 5 Mb ≤ ROH ≤ 10 Mb, 4) ROH ≥ 10 Mb. Groups of clusters are coloured together, and error bars shows 95% confidence intervals. All panels were plotted using the statistical computing language R58 and the package ggplot2.

To complement this non-parametric modelling, we also recorded and compared sharing of IBD-segments and Runs-of-Homozygosity (ROH) within individual NL clusters and Irish or British clusters. We compared the relationship between average number of, and total length of, IBD-segments >3cM and <15cM shared between individuals placed in the same cluster (Figure 4C). NL clusters consistently present higher levels of IBD sharing than British or Irish clusters, and in some cases (e.g., Trinity and Conception Bay clusters) higher than Orkney and Wales. When compared to Orkney or Wales, NL clusters share slightly fewer segments on average even though the total length shared is comparable. This suggests that the increase of relatedness in NL is more recent than Orkney or Wales, or that on average the Ne is overall higher in NL. Elevated IBD levels within NL are also supported by IBD sharing patterns between NL clusters compared to between Irish or British clusters (Figure S4.4), showing a general pattern of elevated haplotype sharing across the province as well as within specific genetic communities. Also supportive of a general elevation of haplotype sharing, ROH levels in NL are also higher than the average in Ireland or Britain (Figure 4D), with some NL clusters in Trinity Bay (for example) exhibiting particularly high levels. This increase in ROH seems driven by longer ROH, consistent with relatively recent isolation.

Finally, we further investigated the evidence of NL-specific genetic drift, to inform on the suitability of NL as an ideal study population where there is an enrichment of rare functional variation. Utilising Patterson’s D statistic37, we first confirmed Irish-English comparative affinities in each NL *fineSTRUCTURE* cluster testing D(YRI, NL; Ireland, England) (Figure 5A), and where an excess of Irish alleles would result in a negative test statistic. We find that the four NL clusters identified with Irish ancestry in our haplotype analysis (Figure 3) are confirmed to present an excess of Irish alleles when compared to English references (Figure 5A). Next, we tested for NL-specific drift by generating two D test statistics for each NL cluster. One arrangement of populations tested the comparative allele shifting between Irish references and 500 random NL individuals not members of the tested cluster, and the other arrangement comparing English references to the same 500 random NL subset (Figure 5B). An excess of NL allele sharing in both tests may indicate NL-specific drift independent of Ireland or England. We find that most NL clusters, but especially Conception Bay S 2 and Conception Bay W, present excess NL drift (Figure 5B). In the case of Conception Bay S 2 and Conception Bay W this is associated with high IBD sharing within those clusters, which would be consistent with an isolated community experiencing excess genetic drift.

**Figure 5.**
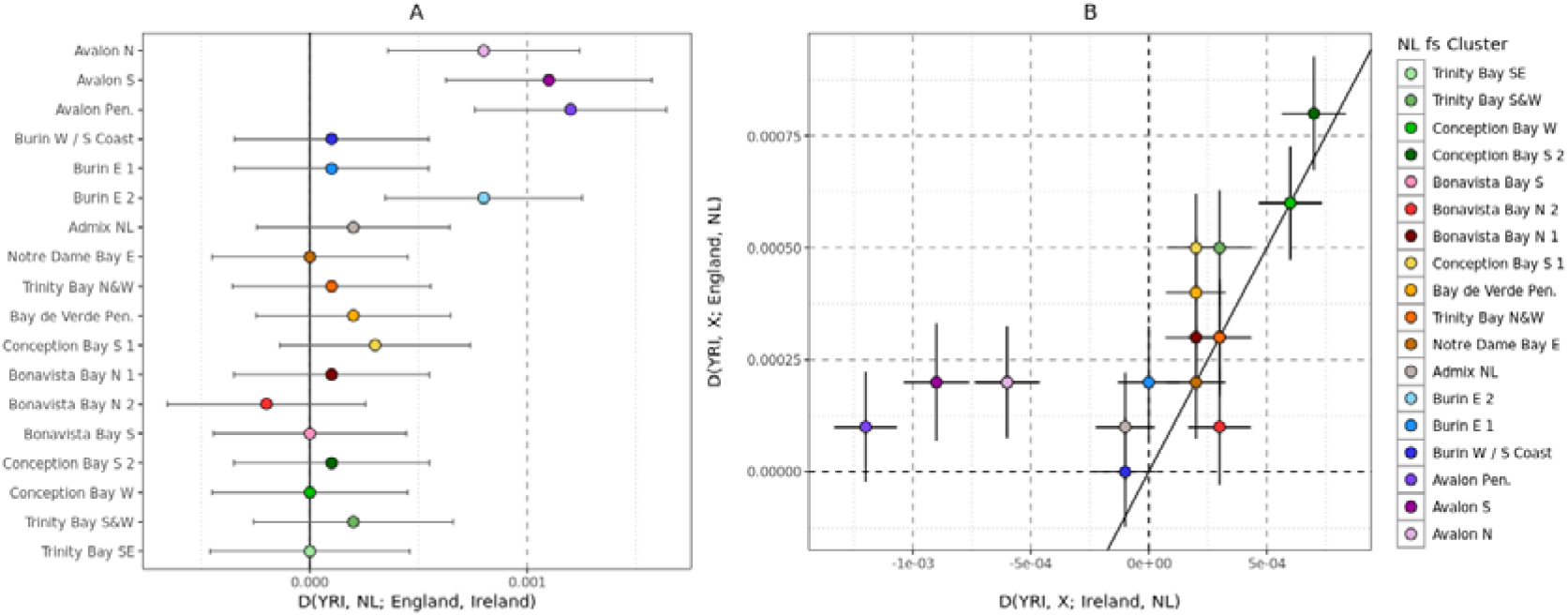
Evidence of NL specific genetic drift from Irish and English source. (A) Patterson’s *D* statistic^37^ testing the allele sharing between NL clusters and English and Irish population references. Positive values indicate excess sharing within NL clusters to Ireland, and negative indicate excess sharing to England. YRI indicates the outgroup for these tests (Yoruban sampled in the 1000 Genomes Project Phase 3). (B) Testing NL-specific drift with Patterson’s *D* statistic. X-axis tests if each NL cluster presents an excess of Irish (negative *D* value) or NL (positive *D* value) alleles, and Y axis tests if each NL cluster presents an excess of English (negative *D* value) or NL (positive *D* value) alleles. All panels were plotted using the statistical computing language R^58^ and the package ggplot2.

## Discussion

In an analysis of 1,807 Newfoundland and Labrador individuals we have elucidated the demographic history of European ancestry within the Canadian province of NL. Irish and British settlers whose ancestry is shared predominantly with the southeast of Ireland and the southwest of England, respectively, underwent a genetic bottleneck approximately 10 to 15 generations ago (300 to 450 years ago assuming generational time of 30 years). This mixture of south-western English and south-eastern Irish ancestry was distributed unevenly across NL, with Irish ancestry predominant in the south and south-east, and English ancestry predominant elsewhere. Post-migration, genetic structure shows geographic stratification around the bays of the island due to population isolation and then expansion. In these populations genetic homogeneity was elevated with increased haplotype sharing both between and within individuals, resulting in NL-specific drift of allele frequencies. The St John’s region of NL in northern Avalon Peninsula is associated with wide-spread haplotype sharing with other regions of NL, suggesting that the urban area was a target of post-settlement migration and resultant admixture from across the province.

Our analysis has substantially expanded our understanding of the genetics of a population with evidence of genetic isolation and bottlenecks. Previous work on rare Mendelian conditions^9-13^ in NL has highlighted the potential of the population for genetic mapping efforts, a case explicitly argued for previously^1^. Separately, previous analysis of 494 NL individuals^3^ showed initial evidence of isolation and genetic structure based along British/Irish ancestry associated with Christian Catholic/Protestant denomination. However, for this study, using a much large NL sample, more genetic markers and new statistical approaches, we are able to extend this work to show in fine detail the genetic structure across the island of Newfoundland. Further building on these observations, our results on the extent of haplotype sharing and genetic homogeneity in NL show that they are comparable to the Orkney Islands, a well-established isolate^38-40^. Evidence of a bottleneck as well as mixing of English and Irish ancestry is remarkably consistent with the dates of major English/Irish settlement within the 18^th^ and 19^th^ centuries (i.e., consistent with estimates of 10 generations ago). In addition, these results have furthered our understanding of European settler genetic history in NL which is of interest to genetic genealogy as well as historical research. Our data indicates that a Protestant or Catholic religious background still acts as a proxy for English or Irish ancestry^33^ even after 300 years of settlement The fact that Irish was predominantly spoken in some of these communities until the early-20^th^ century in Newfoundland^41^ is consistent with these observations. Further, using detailed Irish and British ancestry references, we were able to trace these sources to the specific regions of southeast Ireland and southwest England, greatly improving upon the observations of Zhai et al. This work also highlights the use of detailed reference datasets^21,28^ with geographic or regional data in elucidating human history.

Surprisingly, within our analysis of Britain and Ireland to identify reference clusters for ancestry estimation, we found our IBD-network method was able to detect population structure within England that the *fineSTRUCTURE* method was unable to^20^. This allowed us to genetically differentiate the south-west of England from the rest of England and highlights the potential of relatively simple IBD-based clustering methods that not only scale to large datasets but provide novel insights where other more computationally intensive methods cannot. It is possible that the IBD-based and *ChromoPainter*-based approaches are accessing subtly different time-periods of coalescing genealogies, hence their differing power to detect structure in different regions of Britain. To further highlight this, although IBD-based methods were able to show structure in the south of England, *fineSTRUCTURE* methods were able to differentiate more clusters in north England than our IBD-based approach^20^.

There were some limitations in this work. Without substantial French references, we were unable to explore the degree of French ancestry within NL, where the southern and western coasts of NL have been associated with French migration. In a limited *D* statistic analysis (Figure S4.7) we show with HGDP French reference samples, little evidence of excess French allele sharing in any one NL cluster compared to Ireland or Britain. These observations should be considered as preliminary as this utilised a limited sample size and did not include haplotype data which is more sensitive to sub-continental ancestry differences. Future studies using French Canadian and French samples may further our understanding of French contributions to the NL population structure. Moreover, whilst we were able to group individuals by their religious background, we did not have information on self-reported Irish/British background, limiting our ability to detect any communities of Protestant Irish or Catholic English. Finally, as part of this study we sampled 73 individuals of putative admixed European and Indigenous American ancestry. The original aim of this study was to investigate the genetic history of the European settlers of NL, particularly from Britain and Ireland. While we detected individuals with varying degrees of Indigenous ancestry and included them in analysis of NL fine-scale structure, given the lack of appropriate North American Indigenous references and the small number of participants, we made no further effort to characterise indigenous ancestry in NL. Should a study of the genetic history of the Indigenous Peoples within NL be undertaken in the future, it will require a dedicated study design co-developed with the Indigenous communities, as recommended^42^, to address the priorities of the communities.

In conclusion, we have performed a systematic analysis of the European ancestry within NL. We have highlighted the role of south-western English and south-eastern Irish settlers in forming a genetic landscape with footprints of a substantial bottleneck forming about 10-15 generations ago, consistent with historical records of substantial migration from Ireland and Britain to NL. This bottleneck, and subsequent isolation and expansion of population around the bays of NL, has shaped a genetic landscape with both high differentiation and haplotype sharing within communities. NL’s demographic history has left a unique genetic background, and preliminary data shows that this includes an elevation of normally rare genetic variation to higher frequencies. Coupled with high homogeneity, this genetic profile is ideal for a large population-based genetic mapping efforts, like those achieved in Iceland^43^, Finland^15^, and the Orcadian^40^ populations. Our results not only highlight the use of annotated population genetic references in elucidating human history but also characterise the genetic profile population ideal for human genetic disease mapping in unprecedented detail.

## Methods and Materials

### Datasets

#### Newfoundland and Labrador Cohort

The Newfoundland and Labrador Genome Project (NLGP) is a general population cohort focused on the characterization of Newfoundland and Labrador’s population (Sequence Bio Inc., 2021). The NLGP cohort consists of the initial 2,500 participants for whom medical history and saliva samples were obtained with informed consent under a study protocol approved by the Newfoundland and Labrador Health Research Ethics Board (Ethics Reference #: 2018.243). As part of the participant’s self-reported data, information was collected on their religion and the birthplace of their parental ancestors. Each participant provided a saliva sample using the DNA Genotek Oragene OG-600 collection kit (DNA Genotek, Ottawa, Canada). DNA extracted from these samples was genotyped using the Illumina Global Diversity Array (GDA; Illumina, San Diego, CA). A quality threshold of 99.2% SNP pass rate per sample resulted in 2,446 individuals and 1,721,246 SNPs passing initial genotyping QC. Marker genome positions were made available in both human genome builds 37 and 38.

#### Irish and British References

Irish and British reference genotypes were assembled from the previously reported Irish DNA Atlas^21^, Trinity Student^29^, and the People of the British Isles (PoBI)^28^ datasets. These three datasets were first merged in a combined dataset of 4,469 individuals and 419,033 common SNPs aligned to human genome build 37.

#### World-wide Reference

Genotypes from the combined worldwide references from the 1000 Genomes Project Phase 3 (1KGP3)^44^ and the Human Genome Diversity Project (HGDP)^45^ were accessed from the gnomad v3.1 <u>data release</u>, filtering for autosomal bi-allelic SNPs. A total of 3,942 individuals were used to characterise global patterns of diversity.

### NL-ancestry identification

To characterise global ancestry within the NLGP dataset, we combined the genotypes of 2,446 NL individuals with 3,942 individuals from the world-wide reference dataset. We selected individuals and SNPs with missingness proportions <5%, selecting non-A/T or G/C SNPs with a MAF >2%, and removing SNPs that failed HWE at significance of <1e^-6^. After pruning SNPs in strong LD with the parameters *--indep-pairwise 1000 50 0*.*2* using plink1.9^46,47^, we were left with a dataset of 228,761 common SNPs. We projected the NL individuals onto the principal components calculated from the world-wide references using the *--pca --pca-cluster-names* option and removed ancestry outliers based on their separation with non-European or non-American ancestry clusters, leaving a dataset of 2,417 individuals with either European, Indigenous, or mixed European-Indigenous ancestry.

With the ancestry dataset generated above, we re-extracted SNP genotypes, selecting non-A/T or G/C SNPs with missingness <5%, MAF >2%, and removing SNPs that failed HWE at significance of <1e^-6^. With this filtered set of individuals and genotypes we estimated familial relatedness with KING^48^ and the *--related* option of relatedness estimation – randomly removing one from each pair of individuals with a 3^rd^ degree relationship or closer. We chose a higher threshold than commonly used for population-based analyses as the subsequent haplotype-based analyses have greater sensitivity to relationship between individuals than so-called “unlinked” analyses^20^. The final dataset consisted of 1,807 individuals and 685,221 common SNPs (the “NL_1,807_” dataset).

### NL Population Structure

To perform *ChromoPainter* based haplotype “painting”, we first phased the genotypes into inferred haplotypes using *SHAPEIT* v4^49^ with default parameters, using a recombination map from human genome build 38. Using bcftools^50^ and scripts provided by the authors of *ChromPainter*, we then converted the phased haplotypes from vcf format to the input format required by *ChromoPainter*. With haplotype data encoded in *ChromoPainter* phase and recombrates format, we utilised the *fs* utility which combines *ChromoPainter* and *fineSTRUCTURE* analyses in a single unified pipeline. We estimated the parameters Ne and mu for the calculation of the *ChromoPainter* co-ancestry matrix which records the number of haplotypes that an individual “copies” from other individuals. Using a random subset of chromosomes (3, 9, 16, and 21) we estimated Ne and mu in all NL_1,807_ individuals. With this estimate (otherwise known as stage 1 in *fs*), we calculated the co-ancestry matrix, painting all NL_1,807_ individuals as a mixture of haplotypes donated from every other NL_1,807_ individual (stage 2). For stage 1 and 2, we set the expected haplotype chunks to define a region to 50 rather than the default 100 to reflect the expected homogeneity in the NL cohort^20^. With the resultant “chunk counts” co-ancestry matrix we performed *fineSTRUCTURE* clustering (stage 3), using 2 million burnin and 2 million sampling iterations of *fineSTRUCTURE*’s MCMC clustering algorithm, sampling states every 4,000 iterations. With the MCMC, having sampled the highest posterior probability we performed 200,000 additional tree-building steps (stage 4) to reach a final inferred *maximum a posteriori* (MAP) state and dendrogram of clusters.

We performed several additional analyses to further characterise the clustering of the NL_1,807_ dataset generated by *fineSTRUCTURE*. We processed and visualised the dendrogram in R in part using scripts provided by the authors of *fs*.

We mapped the geographic distribution of clusters by plotting the birthplaces of grandparents born within the same region, so that by restricting the analyses to these grandparents mitigated effects from within-NL migration from the 20^th^ century onward. We first calculated the distance between grandparents of the same NL participant, measured by latitude and longitude. For each grandparent pair (between grandparents *i* and *j*), we calculated the Euclidean distance (Equation 1). Then, calculating the mean distance between all grandparents of the same participant, we filtered for grandparent birthplaces of individuals with a mean grandparental distance ≤ 0.5.

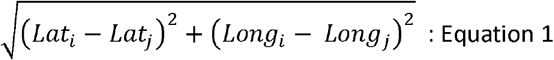

We visualised the multi-dimensional sharing of the *ChromoPainter* co-ancestry matrix by performing principal component analysis using R functions supplied by the authors of *fs*. To visualise the Indigenous ancestry component, and the structure of European ancestry across the province we visualised PC1 versus PC2, and PC2 versus PC3. We estimated genetic distance between clusters using the Hudson F_ST_ estimator^51^ implemented in the <u>admixtools2</u> R package (manuscript in preparation), using the Hudson F_ST_ estimator on a set of 182,222 LD-independent SNPs identified through PLINK using the command --indep-pairwise 1000 50 0.2. We applied the chi-squared test to assess religious association with *fineSTRUCTURE* clusters, computing p-values with simulation because of low counts in the contingency table using the chisq.test function within R. To further enhance the visualisation of the co-ancestry matrix, we applied the t-SNE^32^ and the umap^31^ algorithms implemented in the R packages Rtsne and uwot, respectively. With the exception of setting the t-SNE perplexity to 10, we used default parameters to generate the lower dimensional space.

### NL Irish-British Ancestry

To explore the genetic links between NL, and Ireland and Britain, the NL_1,807_ dataset was combined with the 4,469 individuals of Irish and British ancestry (see “Datasets”). Combining these datasets and selecting SNPs with a genotype missingness <5%, MAF >2% and removing SNPs which failed HWE at significance of <1e^-6^ resulted in a combined dataset with 178,603 common SNPs aligned to the human genome build 37. The combined dataset was then phased together using SHAPEIT v4 to generate a set of inferred NL, Irish, and British haplotypes. The results of a PCA of this dataset is shown in Supplementary Data 3.

To estimate British and Irish ancestry in NL we used sharing of Identical-by-Descent (IBD) segments. We detected segments, > 1cM in length, from phased genotypes using *refinedIBD*^*52*^. We then combined adjacent segments putatively fragments of a true, larger, IBD segment broken up by genotyping or phasing errors with the merge-ibd-segments utility using default parameters.

To explore regional Irish and British ancestry in an expanded sample of Irish and British genotypes (using more Irish genotypes from the Trinity Student dataset than previously analysed^21^) we decided to cluster Irish and British individuals based on genetic sharing. We leveraged the Louvain community detection algorithm^34^ to cluster a network where each individual is a node, and each edge is the total length of IBD in cM that one individual shares with another. We excluded extreme outlier edges which shared >142 cM which are assumed to be more reflective of familial relationships. This approach has been successful in extremely large datasets^53,54^, and was used as it scales to our sample size where other methods such as *fineSTRUCTURE* become a computational burden. The Louvain clustering was performed hierarchically, with three rounds of clustering, each performed on each cluster detected in a previous “level” of clustering. The first level generally split Ireland from Britain, the second level identified broad regions (e.g., NW Ireland, SW Ireland, SE Ireland), and the third level identified the individual clusters reported in Figure 3A. The membership of each third level cluster to each second level group are as follows: CS England (*S&W England, East Anglia, Midlands, Corn. & Devon*), N Britain (*NE England, NW England, NE Scotland, NE Ire & SW Sco*), Wales (*Marches, S Wales, N Wales), Orkney (Orkney*), SW Ireland (*N Kerry, NW Munster, Clare, SW Ireland 1, SW Ireland 2*), SE Ireland (*C Leinster, W Leinster, Leinster, SE Leinster, SW Leinster*), NW Ireland (*Connacht, NW Ulster, N Leinster, N Ire & W Sco*).

To estimate ancestry, we performed a modification of the *nnls* methods reported by Leslie et al^20^, here we leverage IBD-segment sharing as a proxy of recent ancestry. We considered IBD segments shared between NL, Ireland, and Britain that were <15 cM and >3 cM, corresponding to approximately 5 to 15 generations (though this is rough approximation^55^). Our modification of the method generated two matrices of X, and Y. X is a matrix of *n* rows and *m* columns where each row is the total amount of IBD >3cm and <15cM shared between the row individual *n*_*i*_ and each column individual *m*_*j*_. Each row is divided by the sum of IBD summarised in that row so that each *X*_*ij*_ entry is the proportion of IBD recorded in each row. The second matrix is of *m* rows and *m* columns, recording the same for each *Y*_*ij*_ individual-pair. *X* summarises the IBD sharing between *n* “target” individuals and *m* “source” individuals, and *Y* summarises the IBD-sharing between the *m* source individuals and themselves. We use *nnls* to estimate the ancestry proportions explained by these IBD-sharing patterns, modelling each target individual as a mixture of the source individuals.

We performed this modified *nnls* method to estimate sharing proportions in each NL individual as a “target” and leveraged every Irish or British individual as a “source” of IBD sharing. This design modelled NL as a mixture of Irish and British sources, both estimating overall sharing proportions, as well as investigating if certain genetic regions of Ireland and Britain share excess haplotypes with NL, thus allowing the investigation of a specific source of Irish and British ancestry in NL. We used the *cor*.*test()* function provided in R to test the significance and strength of the relationship between the Irish and British ancestry proportions with religious denomination. We tested the correlation between cluster-averaged estimated Irish or English ancestry proportion and the proportion of individuals in that cluster reporting a Catholic or Protestant domination (Anglican or United) background.

To perform supervised *ADMIXTURE*^*30*^ analysis and to calculate Patterson’s *D* statistic^37^ between NL, Irish and British references, and world-wide outgroups we merged the NLGP, Irish and British references, and 1KGP3 and HGDP reference genotypes from the gnomad v3.0 data release together. We selected non-AT/GC SNPs common to all three datasets filtering for SNPs with a missingness < 5% and a MAF > 5%. We additionally removed SNPs in high LD using the plink command –indep-pairwise 1000 50 0.2 for both sets of analyses. In the supervised *ADMIXTURE* analysis, we first applied the *ADMIXTURE* algorithm to just 1KGP3 individuals, setting *k* ancestral populations to five to capture continental ancestry components. We then performed supervised *ADMIXTURE* analysis on the 1,807 NL individuals, using the *P* matrix calculated from 1KGP3 allele frequencies. To estimate *D* statistics between populations we used the *qpDstat* program (version 970) from the admixtools^56^ suite of tools. We defined the “English” reference as British individuals placed within the *S&W England* cluster, and “Irish” reference individuals as either *NW Munster* or *SE Leinster* Irish individuals as these British and Irish clusters contributed the primary signals of the English and Irish ancestry in haplotype analysis.

### NL Haplotype Diversity

We used the IBD-sharing patterns of NL, Ireland, and Britain to estimate recent effective population size (N_e_) over time using the *IBDNe* utility^25^. Using IBD segments > 4cM in length between individuals of the same population label, we estimated N_e_ in Ireland and Britain in the second level IBD clusters (see above). We chose this level as tested groups would have a large enough sample size (> 100 members) for N_e_ estimates to be robust.

To visual haplotype sharing, we considered IBD-segments that were shared between individuals who were placed in the same third level IBD cluster (in the case of Irish or British individuals) or *fineSTRUCTURE* cluster (in the case of NL individuals). For each individual, we averaged the number and total length of IBD that that individual shared with another placed in the same cluster. Then for each cluster we calculated the mean and confidence intervals, and then plotted the relationship between the number of segments shared and the total length in Irish and British clusters, and NL clusters.

Complementing IBD-based data, we used *PLINK* to detect Runs-of-Homozygosity (ROH) in NL, Irish, and British individuals. We calculated ROH using the *--homozyg* option, using a window of at least 50 SNPs, allowing for a maximum of 5 missing calls and 1 heterozygous call in each window, a maximum inverse kb/SNP density of 50, a maximum internal gap of SNPs of 1000 kb, and a minimum ROH size of 1500 kb. We detected five NL individuals with high levels of total ROH (> 200Mb). We excluded these individuals from further ROH based analysis reasoning that they represent outliers with closer-than-average parental relations on-top of the isolated background of NL ancestry. For each *fineSTRUCTURE* or IBD-based cluster, we calculated the mean individual total length of ROH (Mb), divided by the total length of the genome used in ROH detection in Mb to estimate F_ROH_, a genetic proxy of the inbreeding coefficient *F*^*57*^. For each cluster we also calculated the same but for ROH placed within specific length category bins: 1.5 Mb ≤ ROH ≤ 2.5 Mb, 2.5 Mb ≤ ROH ≤ 5 Mb, 5 Mb ≤ ROH ≤ 10 Mb, ROH ≥ 10 Mb.

## Acknowledgements

The authors would like to thank all the participants who consented to participate in the Newfoundland and Labrador Genome Project for enabling this research. The authors wish to thank The Centre for Applied Genomics, The Hospital for Sick Children, Toronto, Canada for the genomic services they provided. This work was supported by the National University of Ireland Post-Doctoral Fellowship in the Sciences and Engineering (to E.G.). This publication has emanated from research supported in part by a research grant from Science Foundation Ireland (SFI) under Grant Number 16/RC/3948 and co-funded under the European Regional Development Fund, by FutureNeuro industry partners, and a SFI Career Development Award under grant number 13/CDA/2223. This study makes use of data generated by the People of the British Isles (PoBI) project. A full list of the investigators who contributed to the generation of this data is available from the relevant PoBI papers.

## Author Contributions

M.S.P, E.G, and G.L.C conceived and designed the project. G.L.C and M.S.P jointly supervised this work. E.G. designed and performed the majority of the bioinformatic analyses. H.Z curated the NL geographic and demographic data, and M.Mc and S.D aided in data visualisation as well as NL data management. S.M. and R.R contributed bioinformatic support. R.A.L, R.E.M.S, G.M, and J.C.S provided project support and comments. S.O, M.Me, A.M.M, L.C.B, and W.B contributed reference data. E.G, G.L.C, M.S.P, and A.L.S wrote the manuscript, and all authors reviewed the manuscript.

## References

1. Rahman, P. et al. The Newfoundland population: a unique resource for genetic investigation of complex diseases. Hum Mol Genet 12 Spec No 2, R167–72 (2003).

2. Mannion, J.J. The peopling of Newfoundland : essays in historical geography., (St John’s : Memorial Univ. of Newfoundland, Newfoundland and Labrador, 1977).

3. Zhai, G. et al. Genetic structure of the Newfoundland and Labrador population: founder effects modulate variability. Eur J Hum Genet 24, 1063–70 (2016).

4. Martin, L.J. et al. The population structure of ten Newfoundland outports. Hum Biol 72, 997–1016 (2000).

5. Martijn, C.A. Early Mikmaq Presence in Southern Newfoundland: An Ethnohistorical Perspective, c.1500-1763. Newfoundland and Labrador Studies. 19(2005).

6. Bear, J.C. et al. Persistent genetic isolation in outport Newfoundland. Am J Med Genet 27, 807–30 (1987).

7. Bear, J.C. et al. Inbreeding in outport Newfoundland. Am J Med Genet 29, 649–60 (1988).

8. Service, S. et al. Magnitude and distribution of linkage disequilibrium in population isolates and implications for genome-wide association studies. Nat Genet 38, 556–60 (2006).

9. Warden, G. et al. A population-based study of hereditary non-polyposis colorectal cancer: evidence of pathologic and genetic heterogeneity. Clin Genet 84, 522–30 (2013).

10. Moore, S.J. et al. Clinical and genetic epidemiology of Bardet-Biedl syndrome in Newfoundland: a 22-year prospective, population-based, cohort study. Am J Med Genet A 132A, 352–60 (2005).

11. Moore, S.J. et al. The clinical and genetic epidemiology of neuronal ceroid lipofuscinosis in Newfoundland. Clin Genet 74, 213–22 (2008).

12. Parfrey, P.S. Autosomal-recessive polycystic kidney disease. Kidney Int 67, 1638–48 (2005).

13. Merner, N.D. et al. Arrhythmogenic right ventricular cardiomyopathy type 5 is a fully penetrant, lethal arrhythmic disorder caused by a missense mutation in the TMEM43 gene. Am J Hum Genet 82, 809–21 (2008).

14. Hatzikotoulas, K., Gilly, A. & Zeggini, E. Using population isolates in genetic association studies. Brief Funct Genomics 13, 371–7 (2014).

15. Locke, A.E. et al. Exome sequencing of Finnish isolates enhances rare-variant association power. Nature 572, 323–328 (2019).

16. Halachev, M. et al. Increased ultra-rare variant load in an isolated Scottish population impacts exonic and regulatory regions. PLoS Genet 15, e1008480 (2019).

17. Gravel, S. et al. Demographic history and rare allele sharing among human populations. Proc Natl Acad Sci U S A 108, 11983–8 (2011).

18. Mathieson, I. & McVean, G. Differential confounding of rare and common variants in spatially structured populations. Nat Genet 44, 243–6 (2012).

19. Lawson, D.J., Hellenthal, G., Myers, S. & Falush, D. Inference of population structure using dense haplotype data. PLoS Genet 8, e1002453 (2012).

20. Leslie, S. et al. The fine-scale genetic structure of the British population. Nature 519, 309–314 (2015).

21. Gilbert, E. et al. The Irish DNA Atlas: Revealing Fine-Scale Population Structure and History within Ireland. Sci Rep 7, 17199 (2017).

22. Gilbert, E. et al. The genetic landscape of Scotland and the Isles. Proc Natl Acad Sci U S A 116, 19064–19070 (2019).

23. Byrne, R.P. et al. Insular Celtic population structure and genomic footprints of migration. PLoS Genet 14, e1007152 (2018).

24. Gilbert, E., Carmi, S., Ennis, S., Wilson, J.F. & Cavalleri, G.L. Genomic insights into the population structure and history of the Irish Travellers. Sci Rep 7, 42187 (2017).

25. Browning, S.R. & Browning, B.L. Accurate Non-parametric Estimation of Recent Effective Population Size from Segments of Identity by Descent. Am J Hum Genet 97, 404–18 (2015).

26. Palamara, P.F., Lencz, T., Darvasi, A. & Pe’er, I. Length distributions of identity by descent reveal fine-scale demographic history. Am J Hum Genet 91, 809–22 (2012).

27. Palamara, P.F. & Pe’er, I. Inference of historical migration rates via haplotype sharing. Bioinformatics 29, i180–8 (2013).

28. Winney, B. et al. People of the British Isles: preliminary analysis of genotypes and surnames in a UK-control population. Eur J Hum Genet 20, 203–10 (2012).

29. Desch, K.C. et al. Linkage analysis identifies a locus for plasma von Willebrand factor undetected by genome-wide association. Proc Natl Acad Sci U S A 110, 588–93 (2013).

30. Alexander, D.H., Novembre, J. & Lange, K. Fast model-based estimation of ancestry in unrelated individuals. Genome Res 19, 1655–64 (2009).

31. McInnes, L. & Healy, J. UMAP: Uniform Manifold Approximation and Projection for Dimension Reduction. ArXiv e-prints (2018).

32. van der Maaten, L.J.P. & Hinton, G.E. Visualizing high-dimensional data using t-SNE. J. Mach. Learn. Res 9, 2579–605 (2008).

33. Zurel, H. et al. Characterization of the Y Chromosome in Newfoundland and Labrador: Evidence of a founder effect. Manuscript in prep.

34. Vincent, D., Blondel, V.D., Jean-Loup Guillaume, J.-L., Lambiotte, R. & Lefebvre, E. Fast unfolding of communities in large networks. J. Stat. Mech. P10008(2008).

35. Byrne, R.P. et al. Dutch population structure across space, time and GWAS design. Nat Commun 11, 4556 (2020).

36. Pankratov, V. et al. Differences in local population history at the finest level: the case of the Estonian population. Eur J Hum Genet 28, 1580–1591 (2020).

37. Reich, D., Thangaraj, K., Patterson, N., Price, A.L. & Singh, L. Reconstructing Indian population history. Nature 461, 489–94 (2009).

38. McQuillan, R. et al. Runs of homozygosity in European populations. Am J Hum Genet 83, 359–72 (2008).

39. McWhirter, R.E., McQuillan, R., Visser, E., Counsell, C. & Wilson, J.F. Genome-wide homozygosity and multiple sclerosis in Orkney and Shetland Islanders. Eur J Hum Genet 20, 198–202 (2012).

40. Xue, Y. et al. Enrichment of low-frequency functional variants revealed by whole-genome sequencing of multiple isolated European populations. Nat Commun 8, 15927 (2017).

41. Newfoundland and Labrador Heritage Website.

42. Claw, K.G. et al. A framework for enhancing ethical genomic research with Indigenous communities. Nat Commun 9, 2957 (2018).

43. Gudbjartsson, D.F. et al. Large-scale whole-genome sequencing of the Icelandic population. Nat Genet 47, 435–44 (2015).

44. Genomes Project, C. et al. An integrated map of genetic variation from 1,092 human genomes. Nature 491, 56–65 (2012).

45. Li, J.Z. et al. Worldwide human relationships inferred from genome-wide patterns of variation. Science 319, 1100–4 (2008).

46. Purcell, S. et al. PLINK: a tool set for whole-genome association and population-based linkage analyses. Am J Hum Genet 81, 559–75 (2007).

47. Chang, C.C. et al. Second-generation PLINK: rising to the challenge of larger and richer datasets. Gigascience 4, 7 (2015).

48. Manichaikul, A. et al. Robust relationship inference in genome-wide association studies. Bioinformatics 26, 2867–73 (2010).

49. Delaneau, O., Zagury, J.F., Robinson, M.R., Marchini, J.L. & Dermitzakis, E.T. Accurate, scalable and integrative haplotype estimation. Nat Commun 10, 5436 (2019).

50. Danecek, P. et al. Twelve years of SAMtools and BCFtools. Gigascience 10(2021).

51. Hudson, R.R., Slatkin, M. & Maddison, W.P. Estimation of levels of gene flow from DNA sequence data. Genetics 132, 583–9 (1992).

52. Browning, B.L. & Browning, S.R. Improving the accuracy and efficiency of identity-by-descent detection in population data. Genetics 194, 459–71 (2013).

53. Han, E. et al. Clustering of 770,000 genomes reveals post-colonial population structure of North America. Nat Commun 8, 14238 (2017).

54. Dai, C.L. et al. Population Histories of the United States Revealed through Fine-Scale Migration and Haplotype Analysis. Am J Hum Genet 106, 371–388 (2020).

55. Al-Asadi, H., Petkova, D., Stephens, M. & Novembre, J. Estimating recent migration and population-size surfaces. PLoS Genet 15, e1007908 (2019).

56. Lazaridis, I. et al. Genomic insights into the origin of farming in the ancient Near East. Nature 536, 419–24 (2016).

57. Clark, D.W. et al. Associations of autozygosity with a broad range of human phenotypes. Nat Commun 10, 4957 (2019).

58. Team., R.C. R: A language and environment for statistical computing. R Foundation for Statistical Computing. (2017).

