## Supplemental Appendix for "Newfoundland and Labrador: A mosaic founder population of an Irish and British diaspora from 300 years ago"

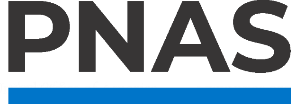

**Supplementary Information for**

Newfoundland and Labrador: A mosaic founder population of an Irish and British diaspora from 300 years ago.

**Authors**: Edmund Gilbert^1,2^, Heather Zurel^3^, Margaret E. MacMillan^3^, Sedat Demiriz^3^, Sadra Mirhendi^3^, Michael Merrigan^4^, Seamus O’Reilly^4^, Anne M. Molloy^5^, Lawrence C. Brody^6^, Walter Bodmer^7^, Richard A. Leach^3^, Roderick E. M. Scott^3^, Gerald Mugford^3^, Ranjit Randhawa^3^, J. Claiborne Stephens^3^, Alison L. Symington^3^, Gianpiero L. Cavalleri^1,2^, Michael S. Phillips^3^.

**Affiliations**:

1. School of Pharmacy and Biomolecular Sciences, Royal College of Surgeons in Ireland, Dublin, Ireland.
2. FutureNeuro SFI Research Centre, Royal College of Surgeons in Ireland, Dublin, Ireland.
3. Sequence Bioinformatics, Inc., St. John’s, Newfoundland, Canada.
4. Genealogical Society of Ireland, Dún Laoghaire, Ireland.
5. School of Medicine, Trinity College, Dublin, Ireland.
6. Genome Technology Branch, National Human Genome Research Institute, National Institutes of Health, Bethesda, MD, 20892, USA.
7. Weatherall Institute of Molecular Medicine, John Radcliffe Hospital, Oxford, United Kingdom.

*** Corresponding Author**: Edmund Gilbert

**This PDF file includes:**

Supplementary Data 1-5.

Figures S1.1 to S5.20.

Tables S1.

SI References.

### **Supplemental Data 1**

#### **NF-Ancestry Identification**

To identify continental-scale ancestry within the NLGP 2,446 individuals we projected the genotypes of the NLGP individuals onto the 3,942 individuals from a combined dataset of individuals from the 1000 Genomes Project (1KGP)^1^ and the Human Genome Diversity Project (HGDP)^2^ – see Methods for details. Shown below are the principal components calculated from this analysis.

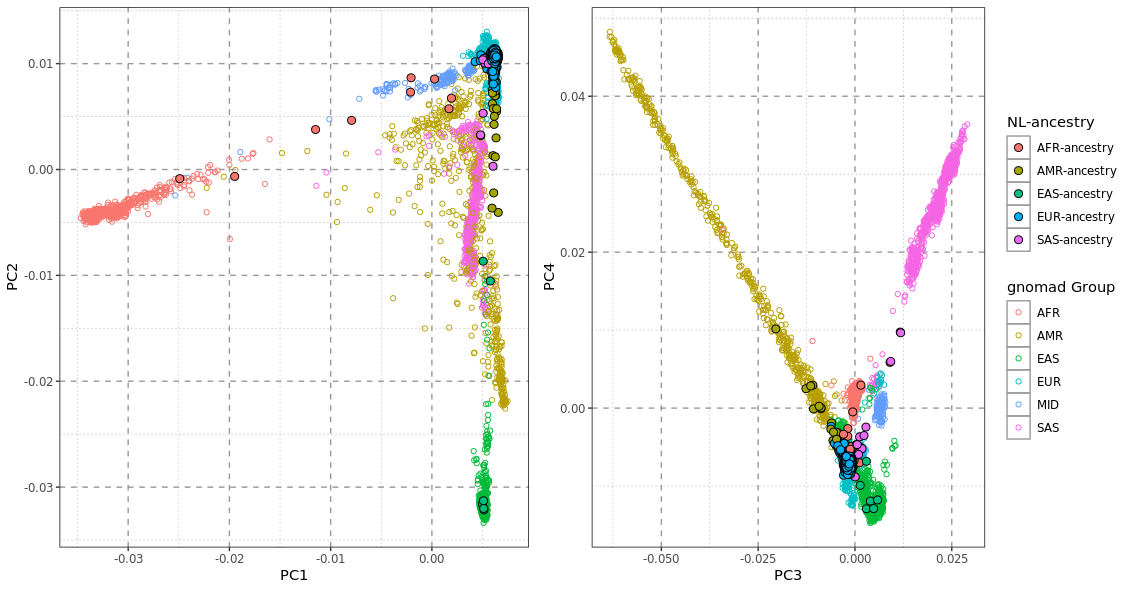

**Figure S1.1** – The first versus second (left) and the third versus fourth (right) principal components (PC) of NLGP individuals (circles with black borders), and ancestry references from either the HGDP or 1KGP datasets (hollow circles). Reference individuals are colour coded by ancestry groups defined by the gnomad dataset; AFR (African), AMR (American), EAS (East Asian), EUR (European), MID (Middle Eastern), SAS (South Asian) NLGP individuals are colour coded by inferred ancestry groups based on PC space.

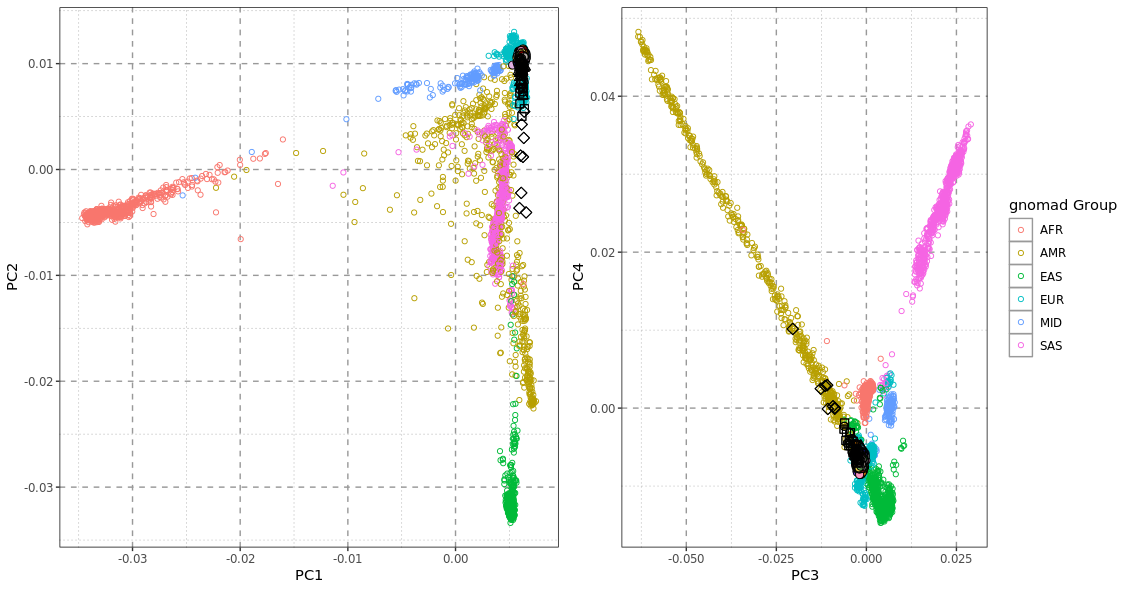

**Figure S1.2** – The first versus second (left) and the third versus fourth (right) principal components (PC) of ancestry references from either the HGDP or 1KGP datasets (hollow circles) with NLGP individuals projected onto the PC space calculated from references. NLGP individuals are only the 1,807 individuals studied as the NL-ancestry dataset, and colour and shape coded according to *fineSTRUCTURE* cluster (see Results). Khaki-colour clusters contain membership of individuals with inferred proportions of Indigenous ancestry.

### **Supplemental Data 2**

#### **NL Geospatial Structure**

Shown below is the geographic distribution of individual NL *fineSTRUCTURE* clusters. Each point represents the birthplace of a grandparent of one of the NL_1,807_ individuals, with a jitter introduced to aid visualisation.

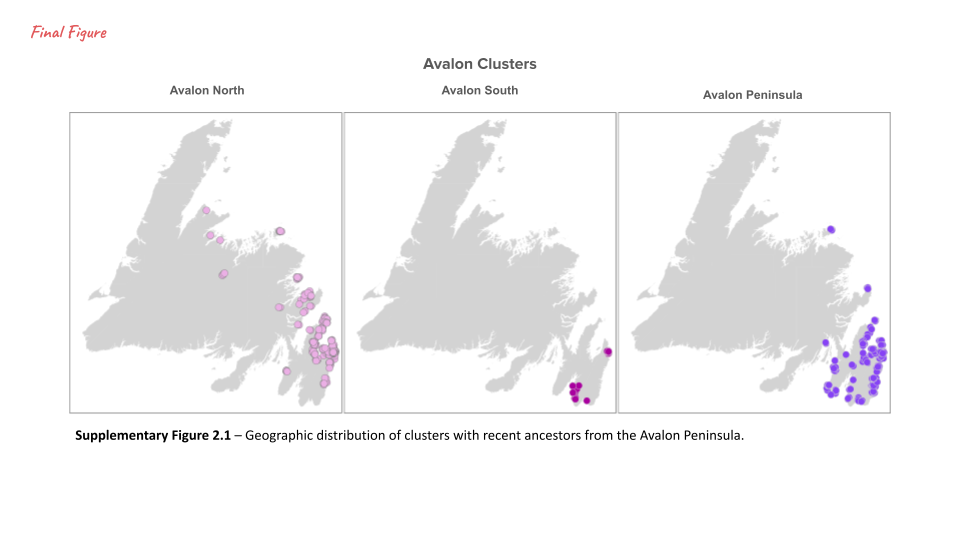

**Figure S2.1** – Geographic distribution of clusters with recent ancestors from the Avalon Peninsula.

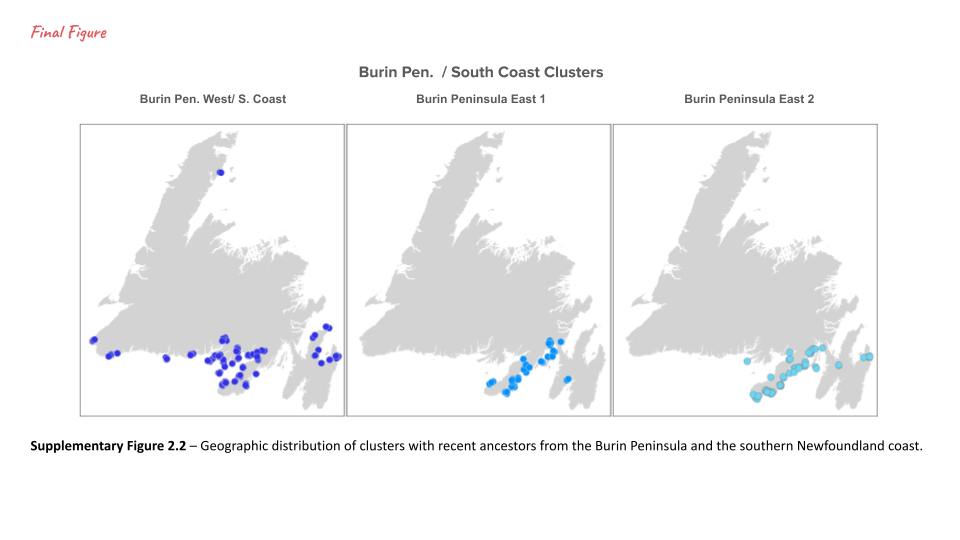

**Figure S2.2** – Geographic distribution of clusters with recent ancestors from the southern Newfoundland coast.

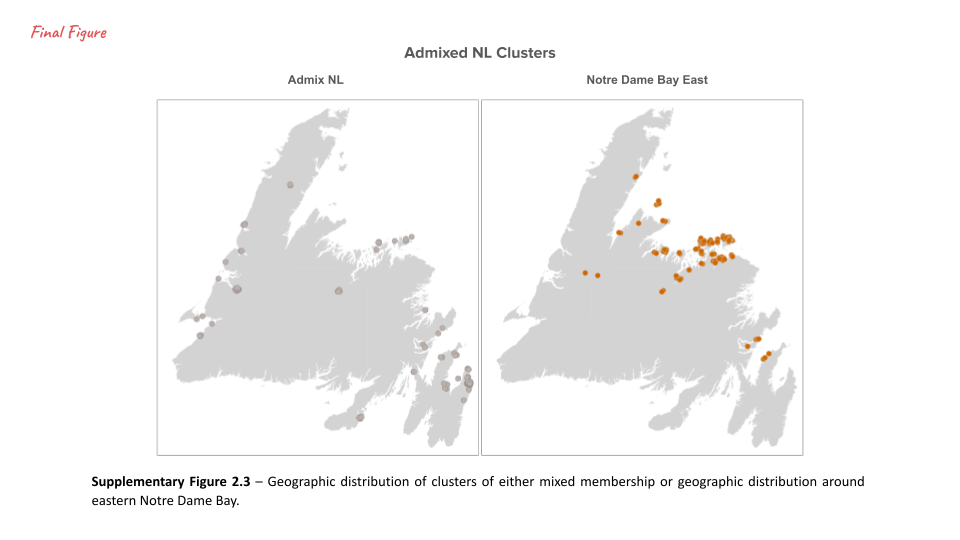

**Figure S2.3** – Geographic distribution of clusters of either mixed membership or geographic distribution around eastern Notre Dame Bay.

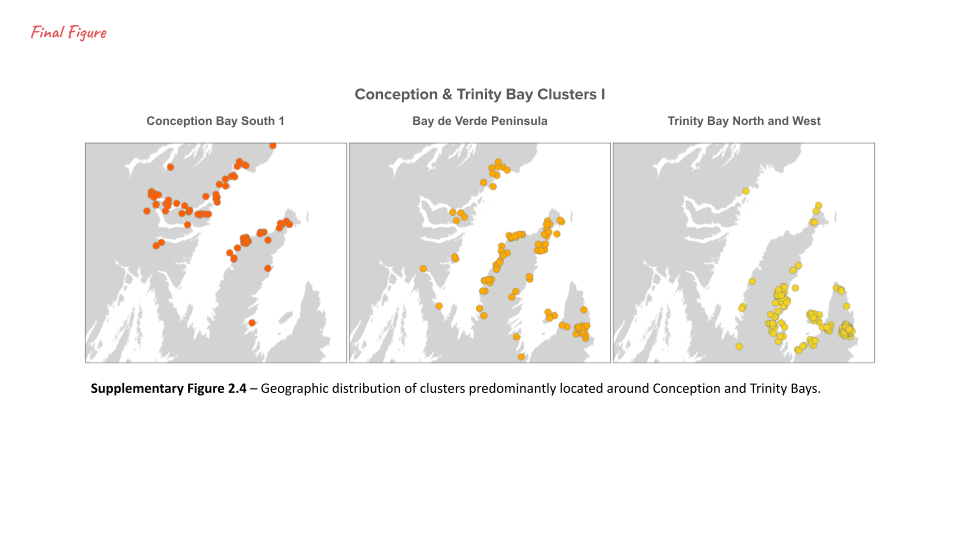

**Figure S2.4** – Geographic distribution of clusters predominantly located around Conception Bay.

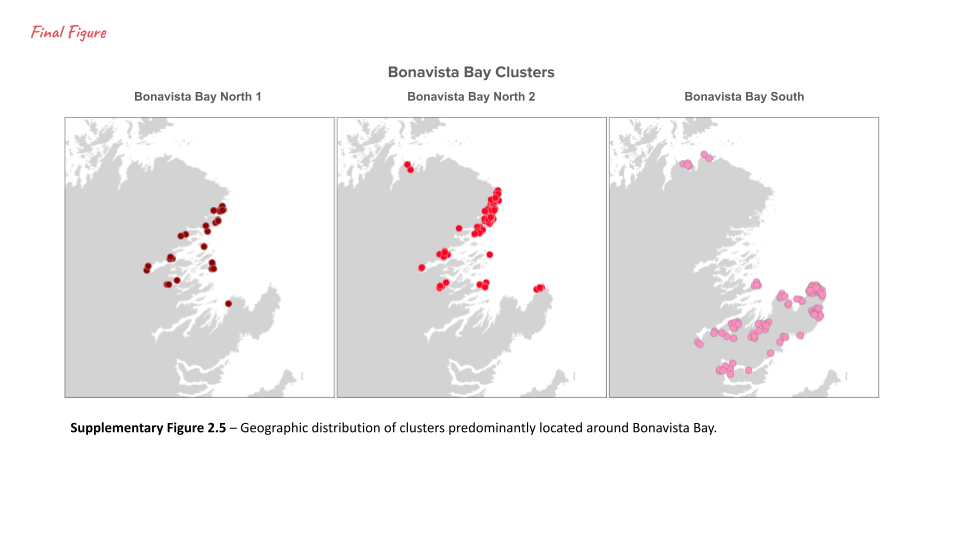

**Figure S2.5** – Geographic distribution of clusters predominantly located around Bonavista Bay.

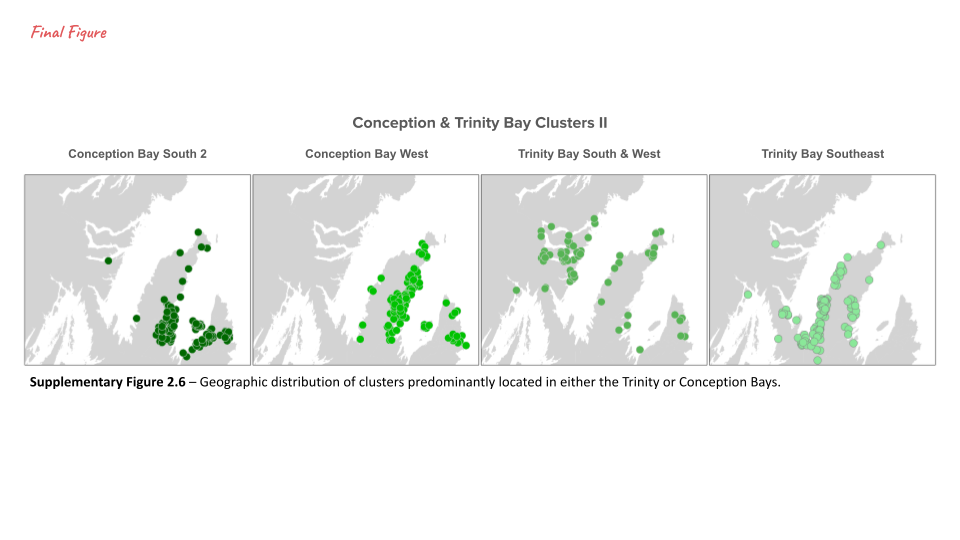

**Figure S2.6** – Geographic distribution of clusters predominantly located in either the Trinity or Conception Bays.

#### **NL Genetic Structure**

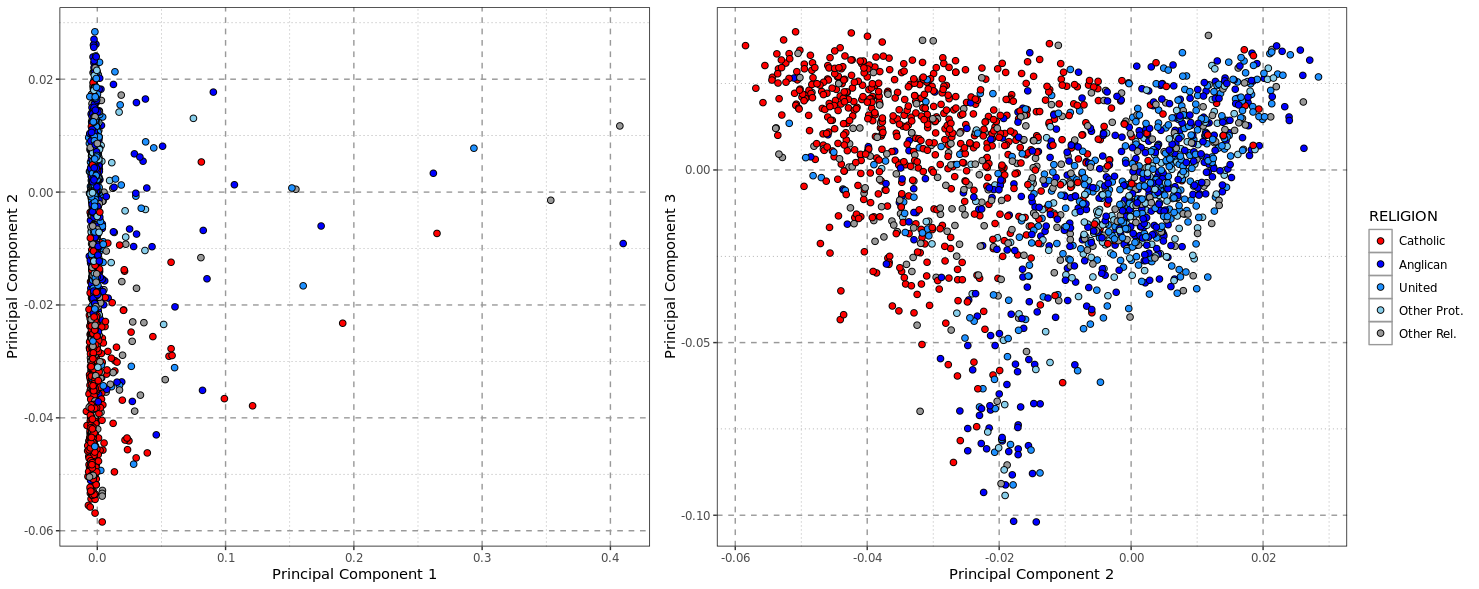

**Figure S2.6** – The first, second, and third principal components calculated from the “chunkcounts” coancestry matrix of the NL_1,807_ individuals. Each point represents the position of one NL individual projected onto the genetic space – colour coded by religious background; with red indicating Catholic, blue indicating Protestant denominations, and grey indicating any other religious background.

#### **NL Indigenous Ancestry**

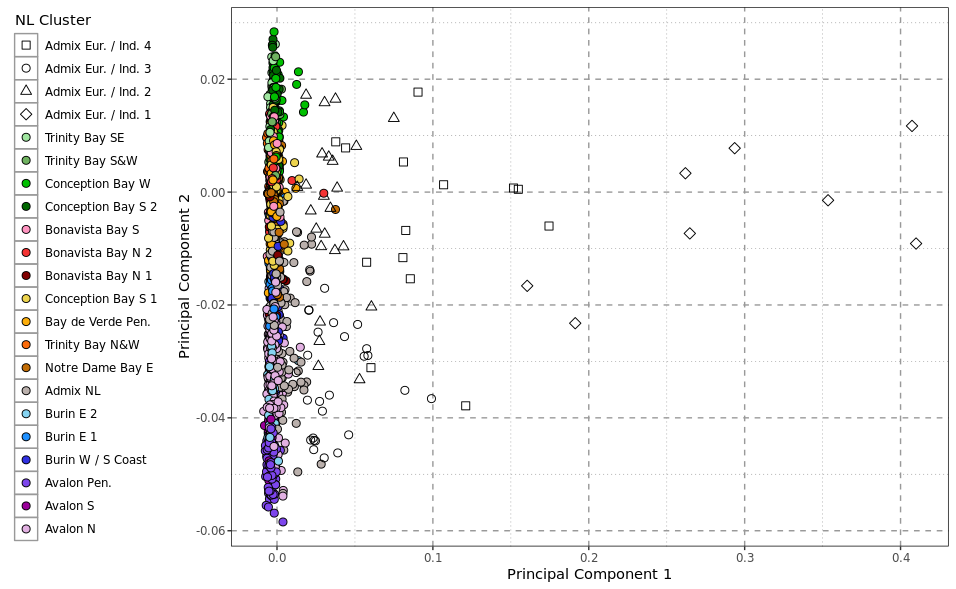

**Figure S2.7** – The first and second principal components calculated from PCA of *ChromoPainter* co-ancestry matrix of 1,807 NL ancestry individuals. Individual genotypes are represented by single points, which are shape and colour coded according to *fineStructure* clustering.

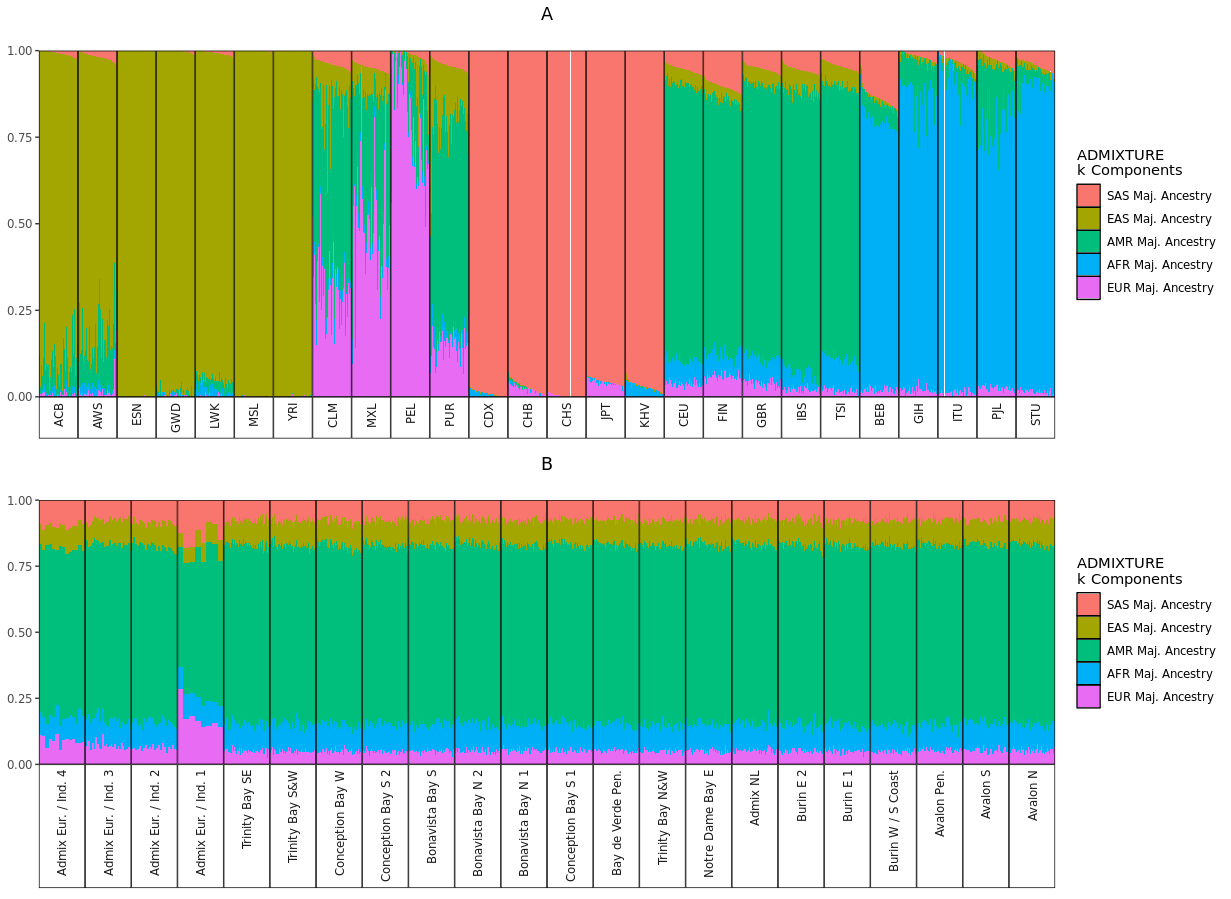

**Figure S2.8** – (A) ADMIXTURE^3^ analysis with *k*=5 ancestry components calculated using allele frequencies of 1000 Genomes Phase 3^4^ individuals, grouped by population and continental meta-group. The five ancestry components are given labels reflective of the continental group that component is maximised in. (B) Supervised ADMIXTURE analysis of 1,807 NL individuals grouped according to fineSTRUCTURE^5^ cluster, where the *k*=5 ancestry components estimated using 1000 Genomes references in A are estimated in the NL individuals.

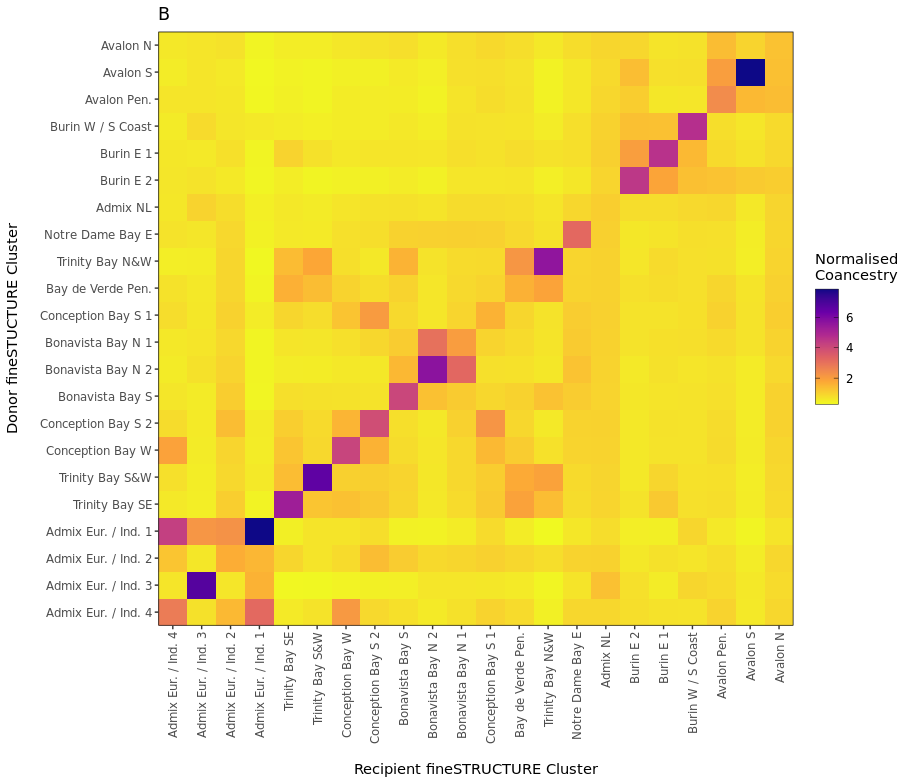

**Figure S2.9** – The column-normalised, cluster-averaged “chunklengths” coancestry matrix showing haplotype sharing between the 22 *fineSTRUCTURE* NL clusters.

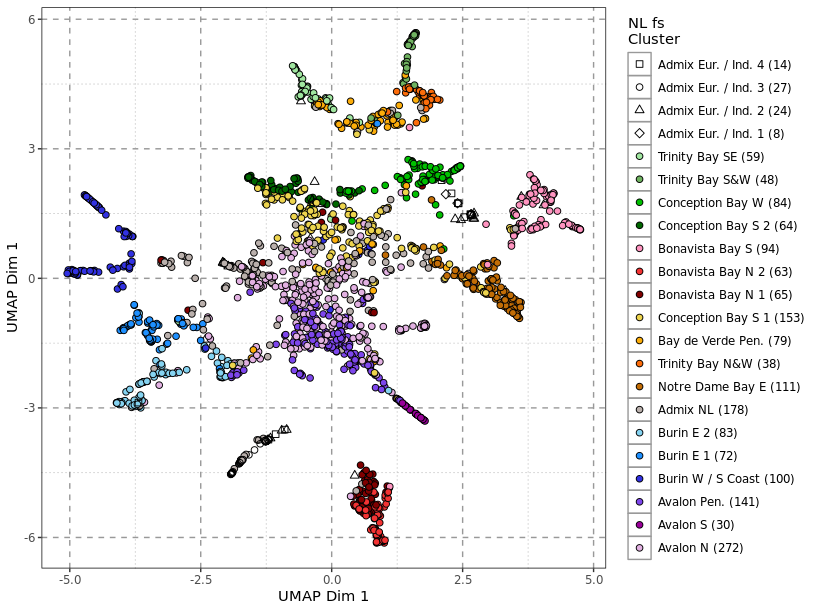

**Figure S2.10** – The umap^6^ projection of the first 20 principal components calculated from the “chunkcounts” coancestry matrix using the R function umap from the uwot package using default parameters. Individual points represent the genetic coordinates of one individual, colour and shape coded according to *fineSTRUCTURE* cluster.

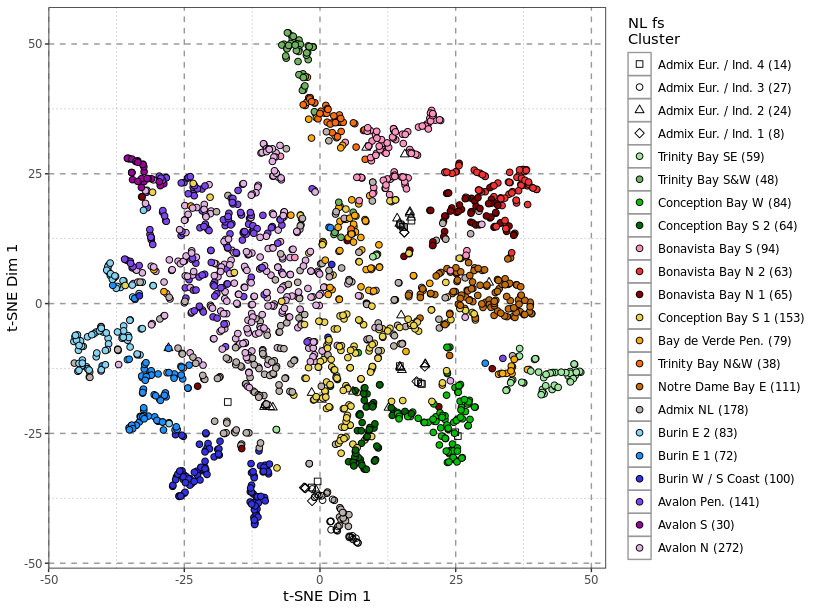

**Figure S2.11** – The t-SNE^7^ projection of the first 20 principal components calculated from the “chunkcounts” coancestry matrix using the R function Rtsne from the Rtsne package using default parameters. Individual points represent the genetic coordinates of one individual, colour and shape coded according to *fineSTRUCTURE* cluster.

### **Supplemental Data 3**

#### **NL and Irish-British Structure**

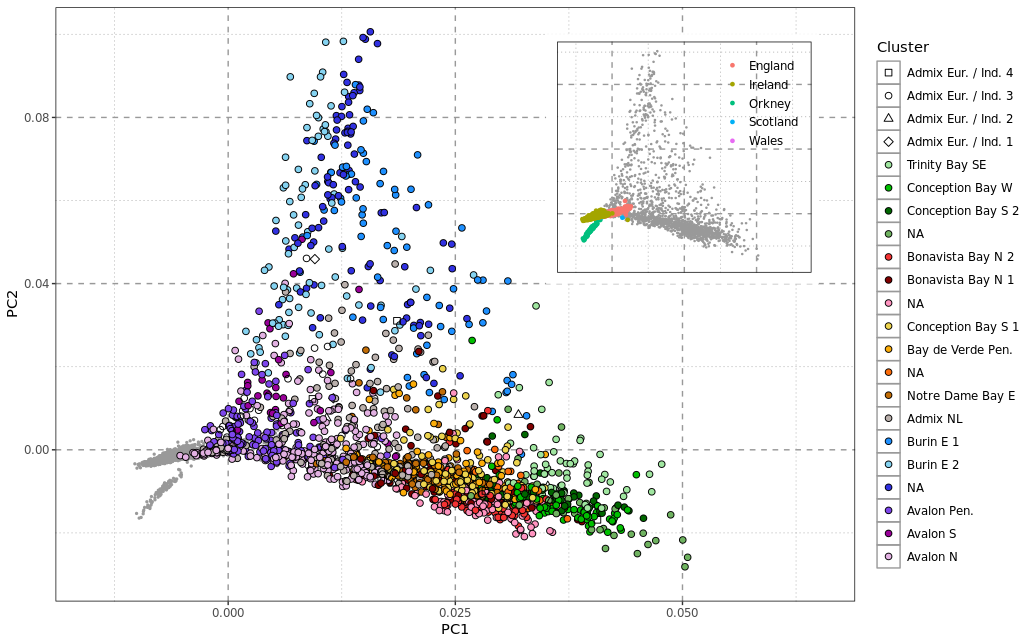

**Figure S3.1** – Principal Component 1 and 2 calculated from co-ancestry matrix of 1,807 NL individuals and 4,469 Irish-British references individuals. NL individuals are colour coded according to *fineSTRUCTURE* cluster membership, Irish or British references are shown as small grey circles with Orkney forming its own cloud. Irish or British individual positions alone are shown as an insert, with yellow indicating Irish, red English, blue Scottish, green Orcadian, and purple Walsh label. The rest of Ireland and Britain forms a cline along principal component 1, with Irish to English as component one increases.

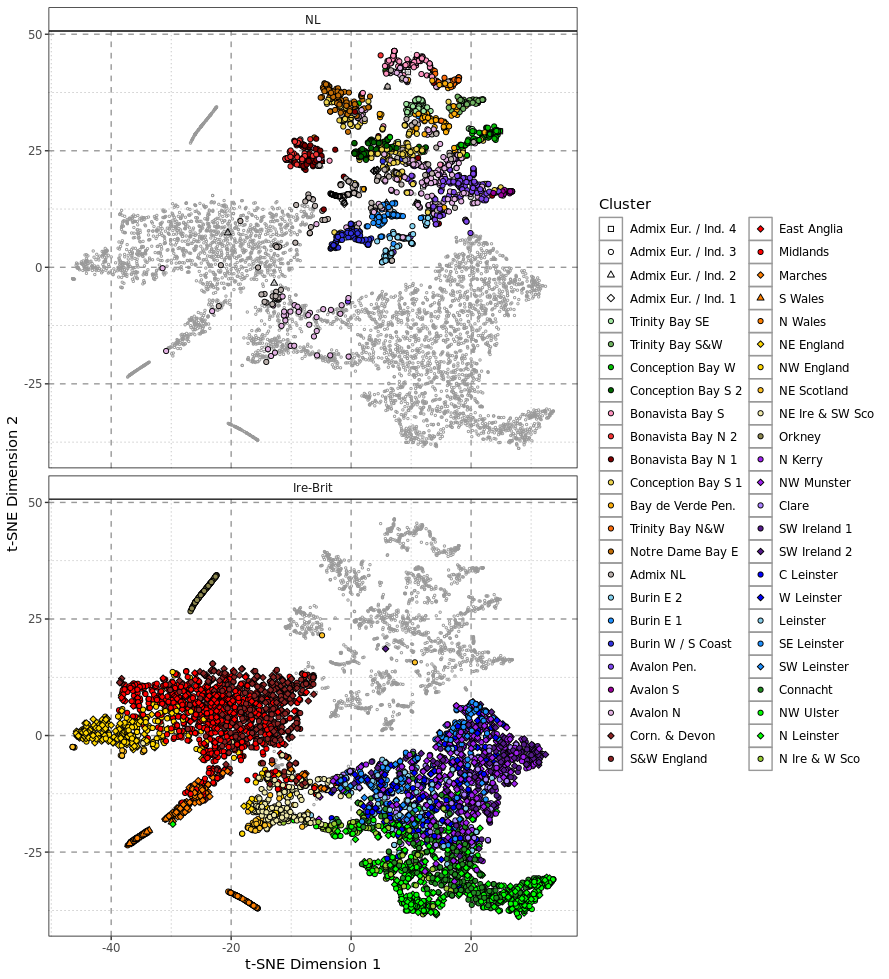

**Figure S3.2** – Genetic coordinates calculated from applying t-SNE to the first 20 principal components calculated from the coancestry matrix of 1,807 NL individuals and 4,469 Irish-British reference individuals. NL and Irish-British references are shown in separate panels, with colour and shape coding indicating *fineSTRUCTURE* (for NL individuals) or IBD-based clusters (for Irish-British individuals).

### **Supplemental Data 4**

#### **Relationship between British-Irish Ancestry and Religious Denomination**

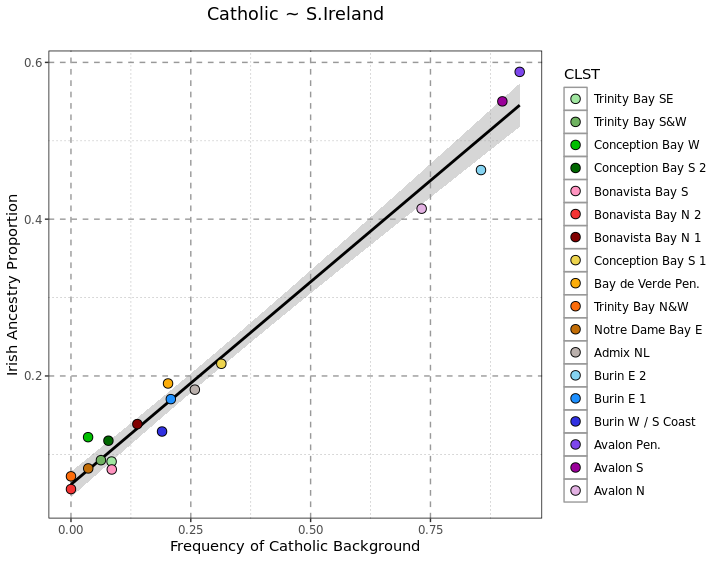

**Figure S4.1** – The per-*fineSTRUCTURE* cluster proportions of Catholic background (x-axis) versus the average estimated Irish ancestry (y-axis). The trend line represents a linear model between the two proportions with 95% confidence shading.

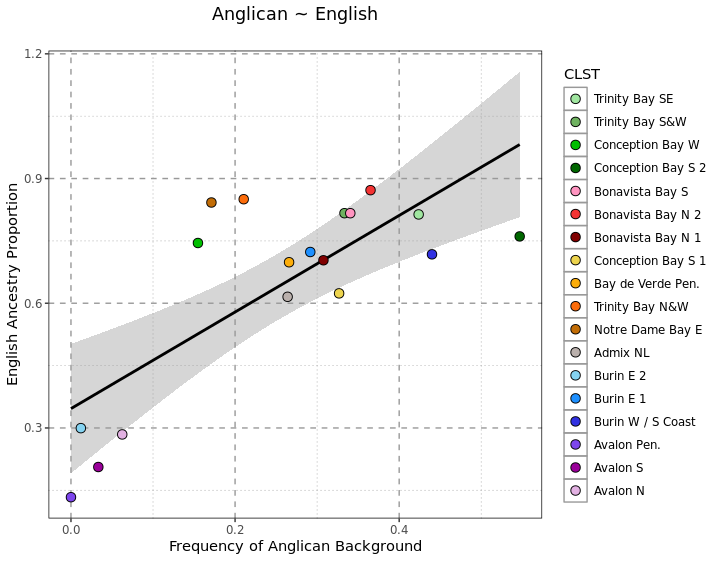

**Figure S4.2** – The per-*fineSTRUCTURE* cluster proportions of Anglican background (x-axis) versus the average estimated English ancestry (y-axis). The trend line represents a linear model between the two proportions with 95% confidence shading.

#### **IBD Copying Profiles Across Ireland, Britain, and NL**

In addition to modelling each NL *fineSTRUCTURE* cluster as a mixture of Irish or British haplotypes we generated broad ancestry profiles in an “unsupervised” analysis, investigating overall IBD sharing patterns between NL, Irish, and British clusters. This analysis would detect both within and without sharing between these three populations. Using the methodology set out in Methods, for each *fineSTRUCTURE* or IBD cluster we recorded IBD segments shared between individuals within that cluster and every other *fineSTRUCTURE* or IBD cluster.

We treated each NL, Irish, and British cluster as a target cluster separately, using every other NL, Irish, or British cluster as a source (Supplementary Figure 4.1). Visualising these pairwise comparisons of target versus source clusters as a heatmap of average ancestry contributions, relationships between groups of clusters become clear. Each of the three broad population groups (NL, Britain, and Ireland) present evidence of genetic regions of inter-related clustering. The patterns in Ireland and Britain are consistent with previous results^8-10^. Within NL, clusters from the Avalon Peninsula region are distinct from the other NL clusters, in agreement with *fineSTRUCTURE* dendrogram ordering. Interestingly, despite high levels of NL-NL IBD sharing patterns, we detected low IBD-sharing in some NL target clusters from British and Irish source clusters. In Avalon for example, this is predominantly from Irish sources, not British. This agrees with the high levels of estimated Irish haplotypes in these NL regions summarised in Figure 3.

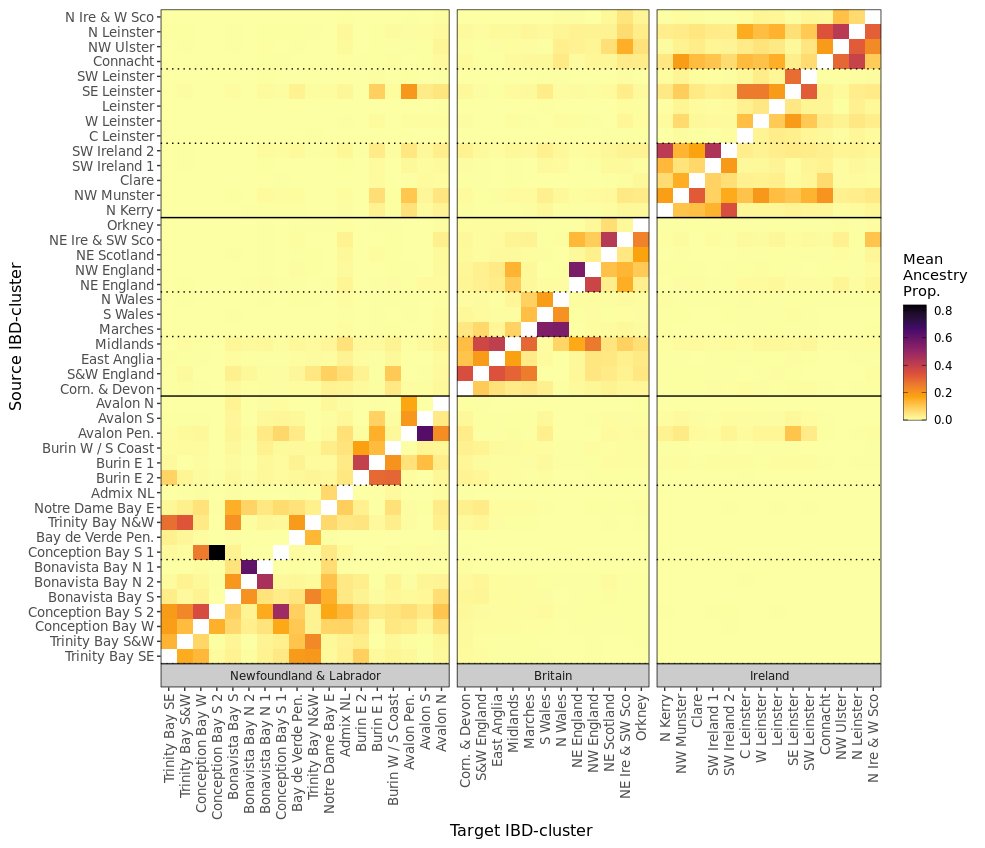

**Figure S4.3** – The mean per-cluster pair “ancestry proportion” matrix calculated using IBD-sharing proportions and the nnls-method^8^ between all NL, Irish, and British cluster pairs. Solid horizontal lines indicate groups of overall populations, and dotted horizontal lines indicate broad sub-groups of clusters within those populations.

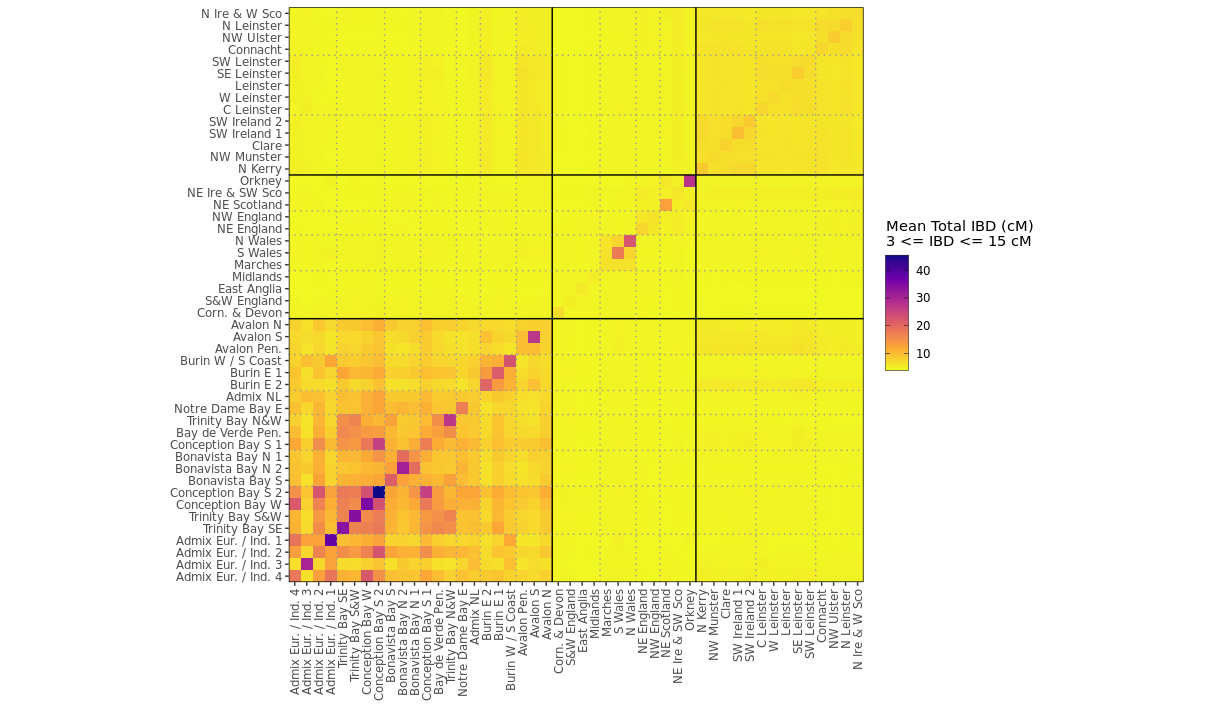

**Figure S4.4** – The average total length of IBD > 3cM and < 15 cM between all NL and Irish and British clusters. Clusters are grouped according to NL, British, or Irish membership. These groups are divided by solid black lines, and groups of clusters within NL, Ireland, or Britain are shown with dotted grey lines.

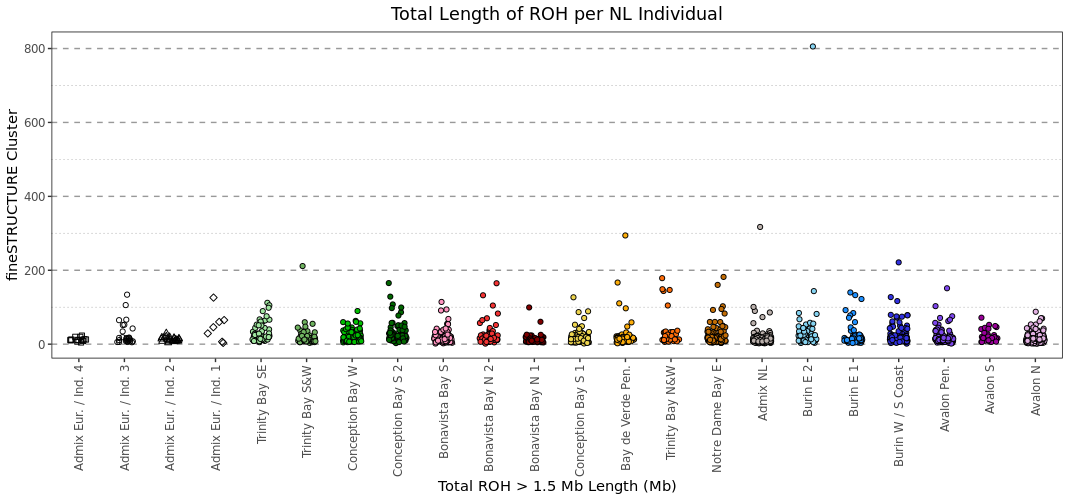

**Figure S4.5** – Total length of Runs of Homozygosity per NL individual (Mb), grouped according to *fineSTRUCTURE* cluster membership.

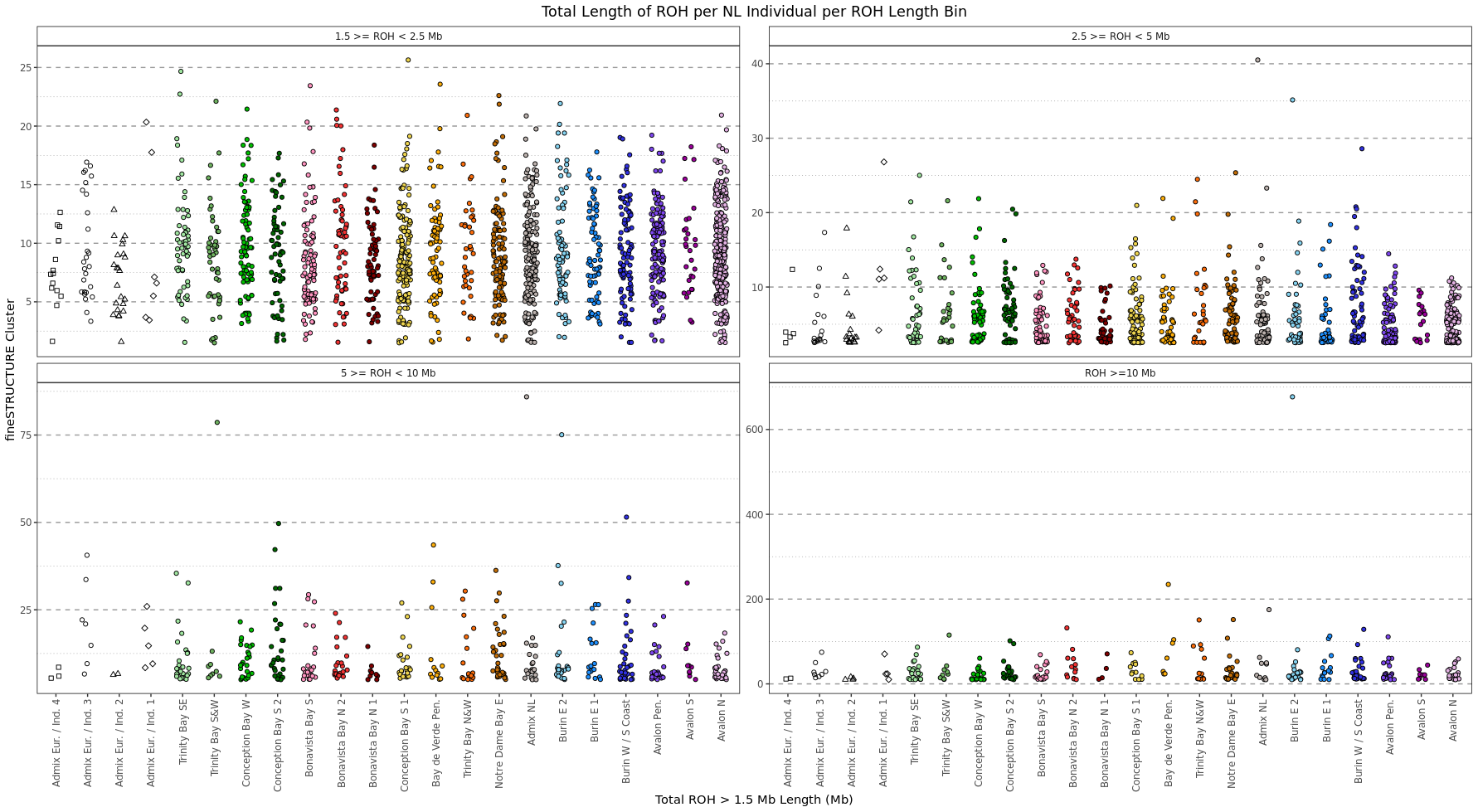

**Figure S4.6** - Total length of Runs of Homozygosity per NL individual (Mb), grouped according to *fineSTRUCTURE* cluster membership, and show per ROH length category bin.

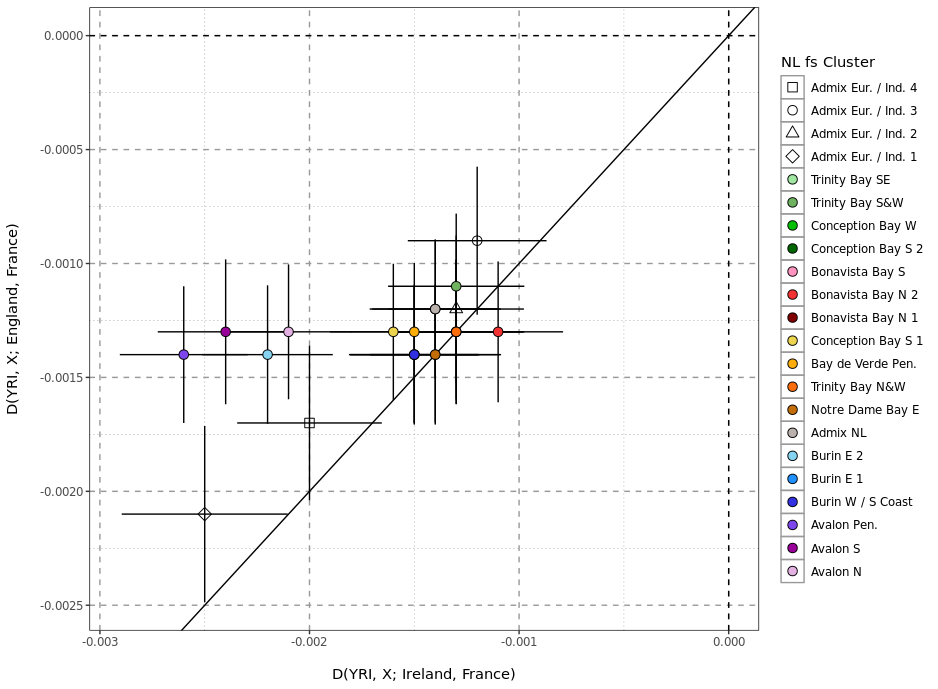

**Figure S4.7** – Patterson’s *D* statistic analysis of Irish-Enlish-French ancestry, where negative values on the x and y axes indicate excess allele sharing between NL and Irish or English sources (respectively), and positive values on the x or y axes indicate excess allele sharing between NL and French sources.

### **Supplemental Data 5**

#### **Evidence of Irish-British Admixture in NL from fastGLOBETROTTER**

To complement the IBD-based ancestry contribution method reported in our main results we applied the *fastGLOBETROTTER* method to the joint Irish-British and NL dataset. *fastGLOBETROTTER* is an optimised extension of *GLOBETROTTER*^11^, providing faster and more accurate inferences of admixture events. We utilised to *fastGLOBETROTTER* to provide evidence of a mixture of Irish and British haplotypes forming the modern European ancestry across NL. We specifically tested the evidence of detectable admixture events between Irish or British source clusters in any NL *fineSTRUCTURE* cluster, and the mixture content of any detected admixture events.

We tested each NL *fineSTRUCTURE* cluster for evidence of a “single-event” admixture signal using the Irish and British IBD-clusters as surrogates for the sources of any detectable event. Using phased haplotype data previously used to detect IBD segments, we generated a *ChromoPainter* co-ancestry “chunklengths” matrix. We “painted” each NL, Irish, or British haplotype as a mixture of every other NL, Irish, or British haplotype using stages 1 and 2 of the *fs* utility^5^. Using the same haplotype data, we generated “painting samples” which are derived samples of the *ChromoPainter* model where each NL individual haplotype was modelled as a mixture of Irish or British haplotypes at each SNP. Leveraging this copying “samples” data, and the total co-ancestry matrix, we performed *fastGLOBETROTTER* analysis, using fastGT mode 1 with 100 bootstraps of the estimated admixture date. We focused on detected “single-event” admixture which had >95% of bootstrap dates replicated within the bounds of accurate *fastGLOBETROTTER* estimation, between 1 and 200 generations ago. We also focussed analysis on events where the admixture model fit well with the underlying data, restricting to event with a maximal R2 goodness-of-fit for the inferred admixture of > 0.5.

As well as estimating admixture in each individual NL *fineSTRUCTURE* cluster, we estimated evidence of admixture using an “all NL” target, which included grouping all NL individuals except for members of an inferred Indigenous-mixed cluster as one NL-meta-cluster. We also did not estimate such admixture in each separate inferred Indigenous-mixed NL cluster as this did not fit the inferred demographic history of these four clusters. We report the co-ancestry curves (Supplementary Figures 5.2-20), showing the *fastGLOBETOTTER* model fit to the observed data for each of the tested 19 clusters of NL individuals.

We detect significant evidence of one-date admixture events in 3 of the 18 NL clusters (Supplementary Figure 5.1). The average date of these admixture events is 9.5 generations ago, ranging from 9.79 generations in *Avalon N* to 8.59 generations in *Avalon Pen*. Inferred admixture events with the greatest fit to the data (including the results of the “whole NL” analysis) tend to date to 10 generations ago. These tend to predate the historical records of 18^th^ and 19^th^ century settlement, though the *GLOBETROTTER* algorithm’s date estimates can be considered as an upper estimate^8^, so could be viewed as consistent with the historical record. This analysis also further supports our IBD-based results from *nnls* analysis, where the mixing sources are from the south-east of Ireland (with NW Munster acting as the major surrogate for an Irish source) and the south-west of England (with S&W England acting as the major surrogate for an English source). Together, these results support European ancestry in NL as a mixture of ancestries from the southeast of Ireland and the southwest of England – dating to approximately 10 generations ago.

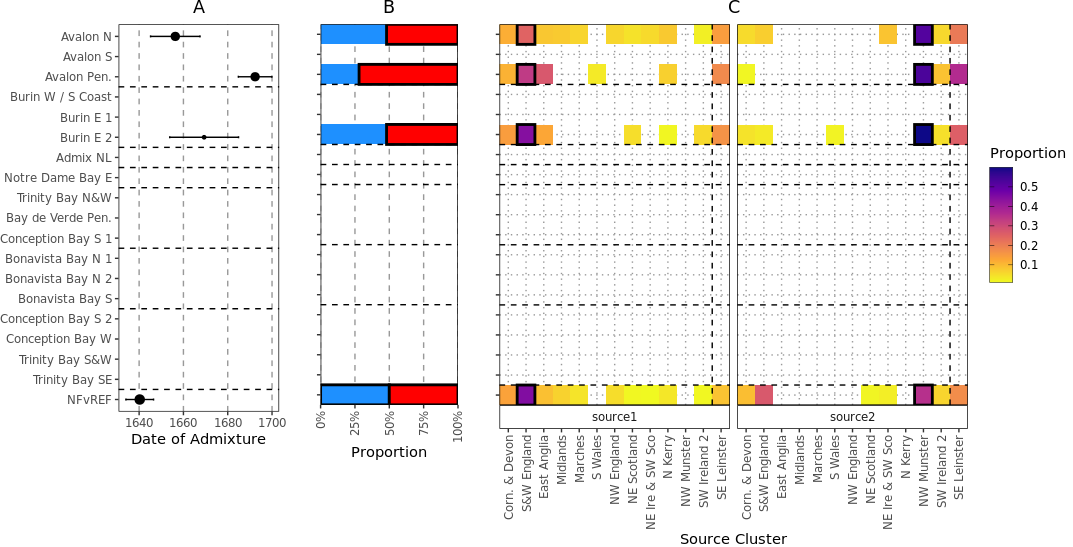

**Figure S5.1 – History of European admixture within Newfoundland and Labrador.** (A) Dates of admixture events estimated by *fastGLOBETROTTER* to be a “one-date” event, and where 95% of the estimated dates from bootstrap replicates where >1 or <200 generations ago, and the maximal R2 goodness-of-fit of the inferred admixture event is > 0.5. Error bars show 95% confidence intervals, and the size of point is proportional to the maximal R^2^ goodness-of-fit of the inferred admixture event to co-ancestry curves. (B) Inferred proportions of the two mixing sources in each significant estimated one-date admixture event. Blue shows “source1” and red “source2”, with a black border showing the majority mixing source. (C) The inferred surrogate sources for source1 and source2, with colour indicating proportion that the surrogate contributes to mixing sources 1 or 2. Black border shows the surrogate which contributes the majority to each mixing source. All panels were plotted using the statistical computing language R^12^ and the package ggplot2.

#### ***fastGLOBETROTTER* Coancestry Curves**

Shown below are the co-ancestry curves constructed by *fastGLOBETROTTER* to infer admixture times and the makeup of mixing groups from a list of surrogates. For each NL *fineSTRUCTURE* we show the curves between the surrogates which contributes the majority to each admixing source (highlighted in Supplementary Figure 5.1). Each curve shows the weighted probability that two positions separated by distance X-axis copy from the pair of populations listed above each panel – with red and green lines showing the *fastGLOBETROTTER* inferred fit for the date estimate (red) or mixing source make-up (green).

**Figure S5.2 – NLvREF**

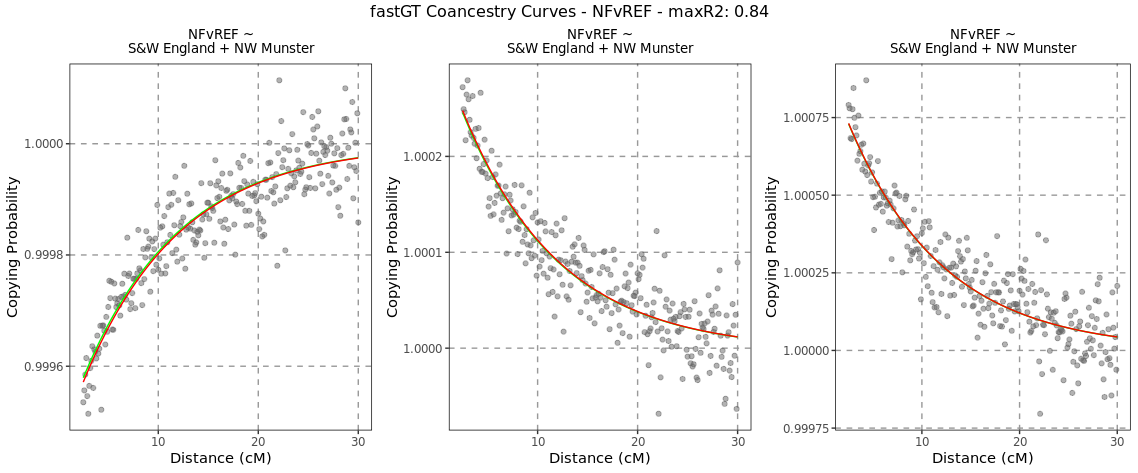

**Figure S5.3 – Avalon N**

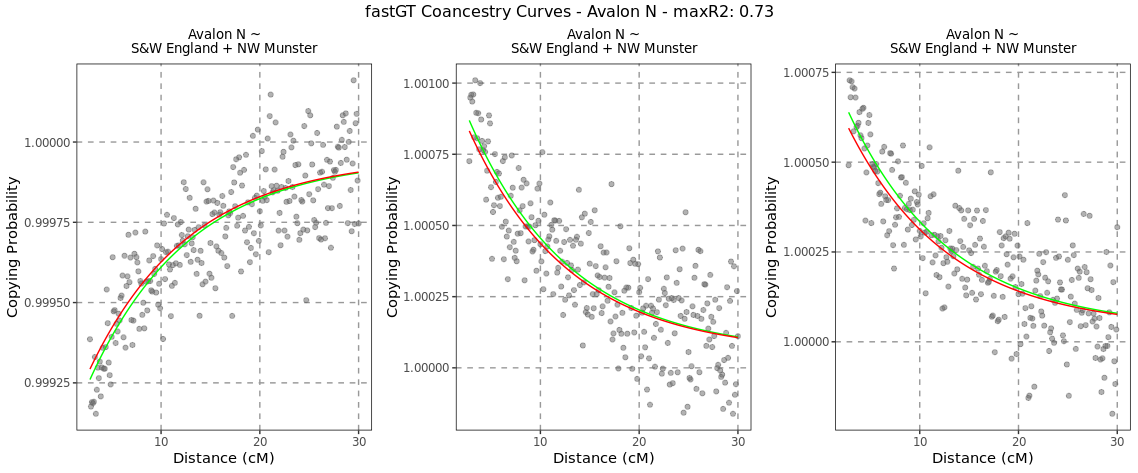

**Figure S5.4 – Avalon S**

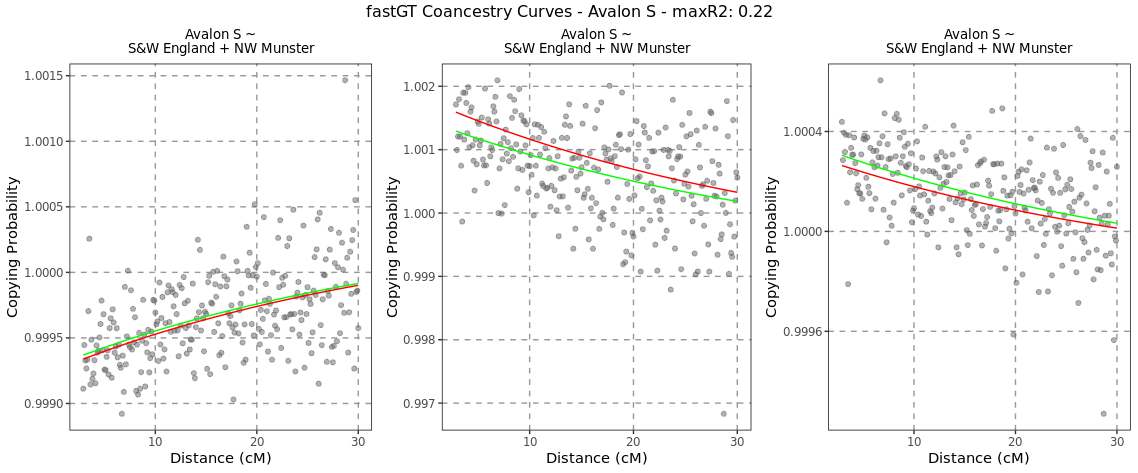

**Figure S5.5 – Avalon Pen.**

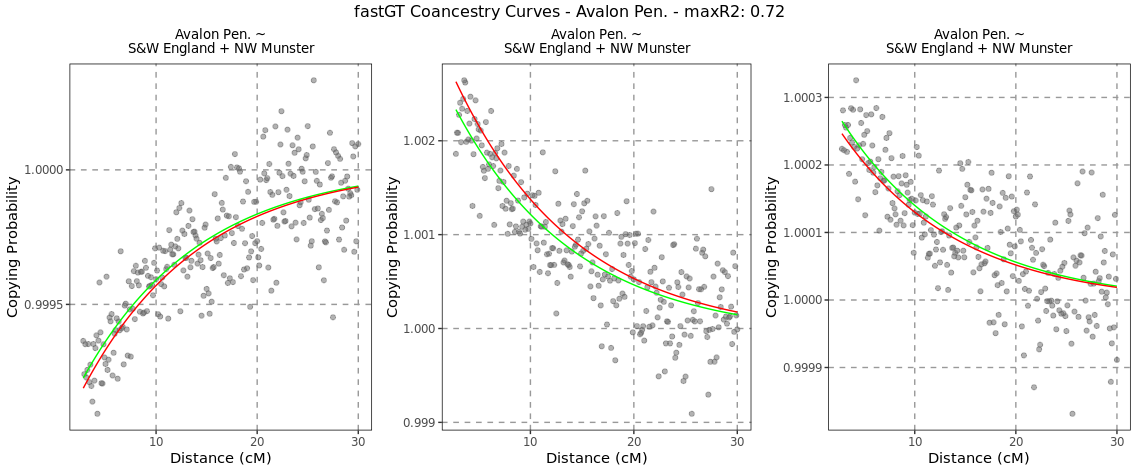

**Figure S5.6 – Burin W/S Coast**

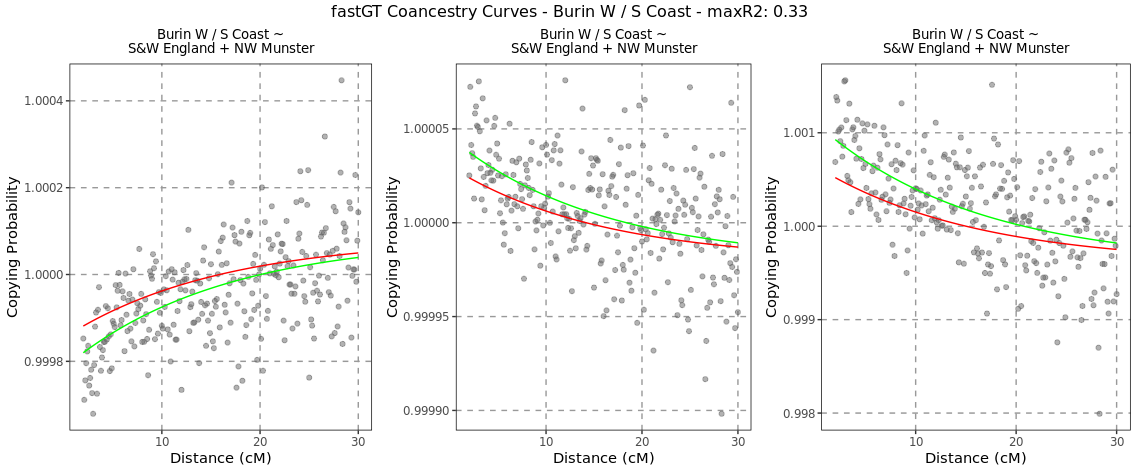

**Figure S5.7 – Burin E 1**

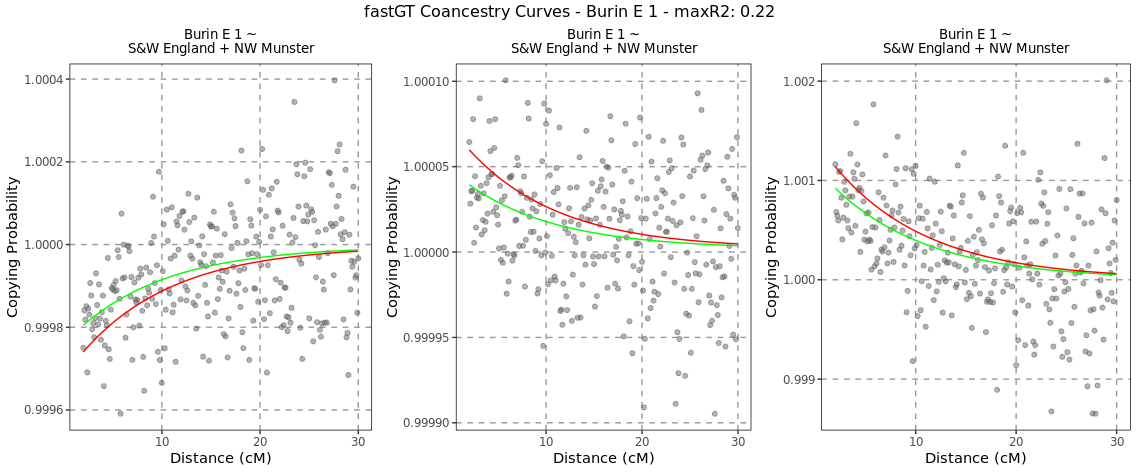

**Figure S5.8 – Burin E 2**

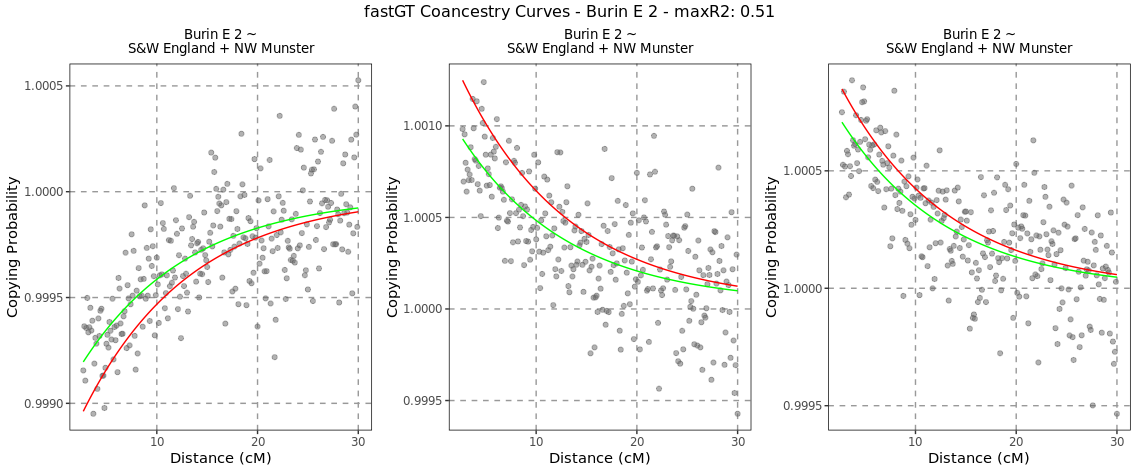

**Figure S5.9 – Admix NL**

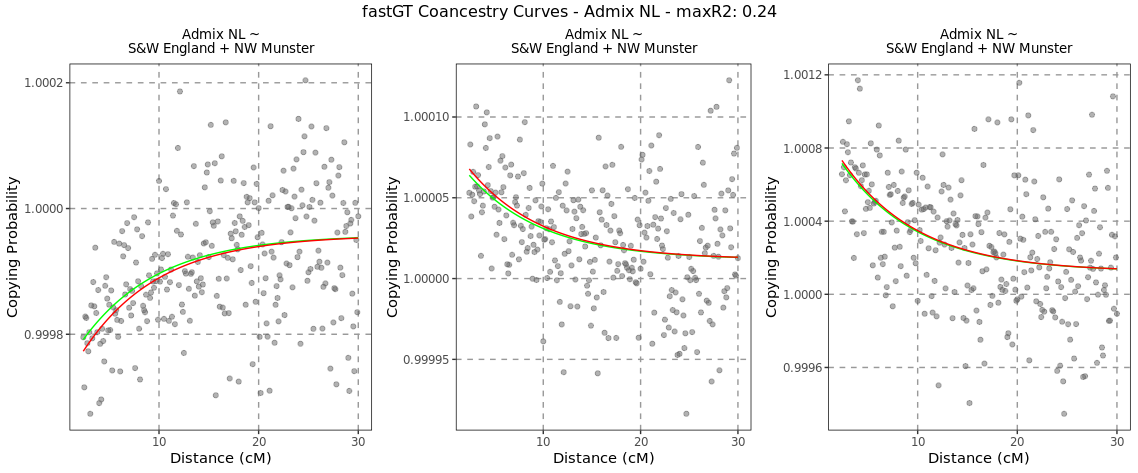

**Figure S5.10 – Notre Dame Bay E**

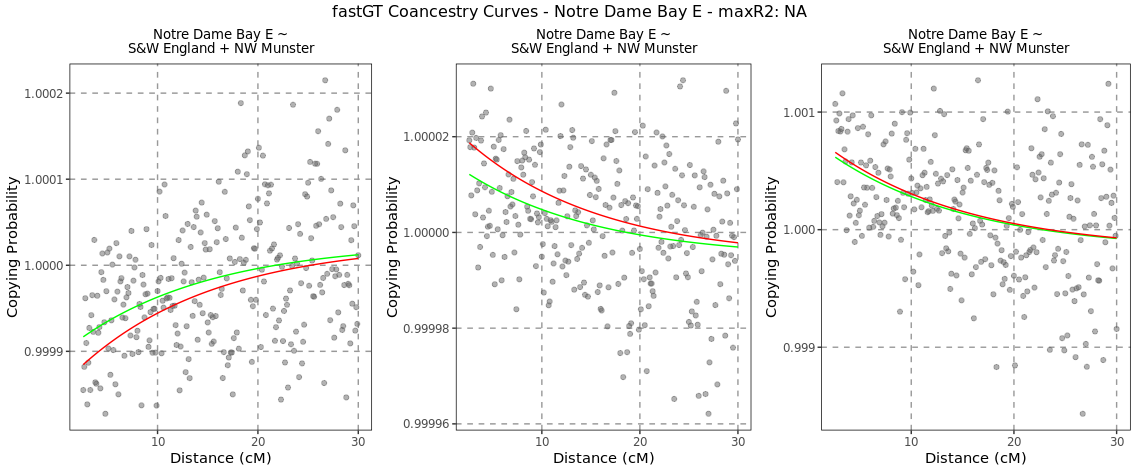

**Figure S5.11 – Trinity Bay N&W**

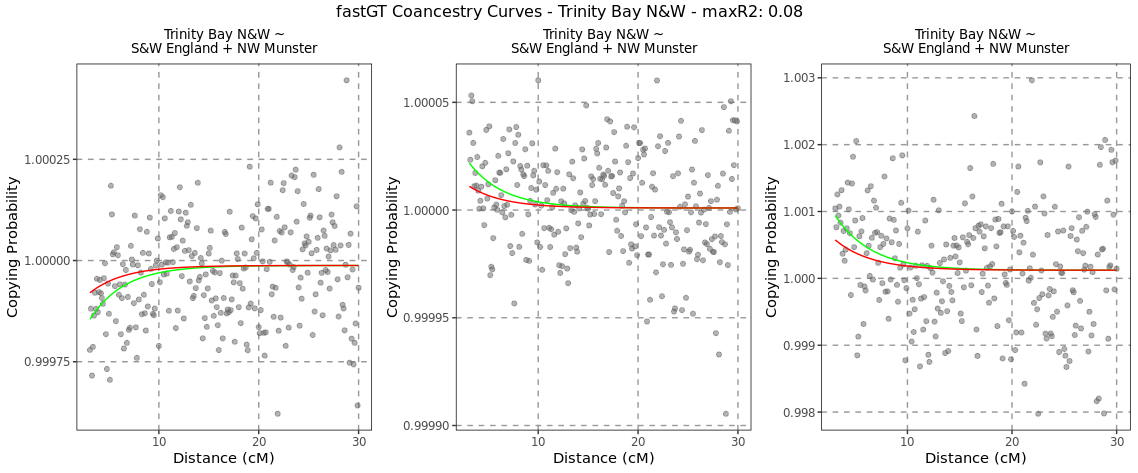

**Figure S5.12 – Bay de Verde Pen.**

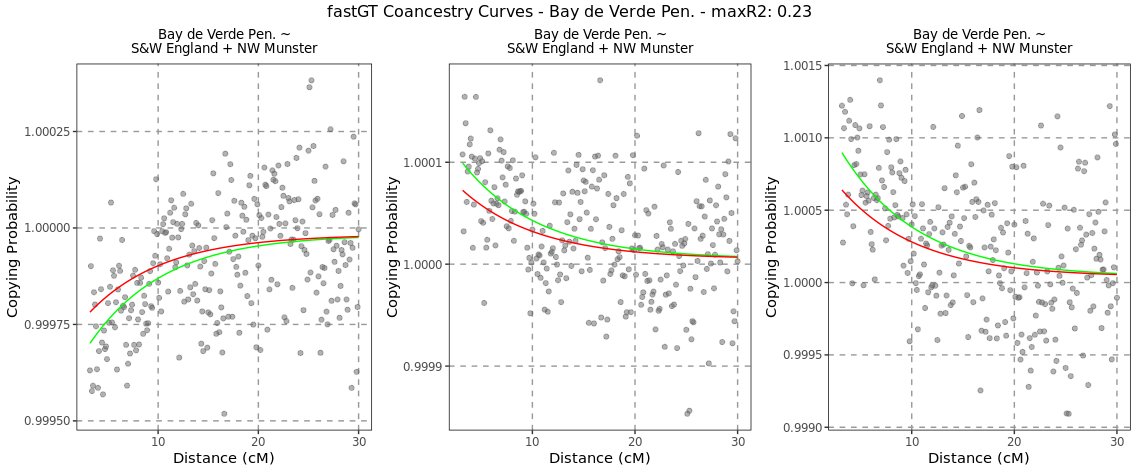

**Figure S5.13 – Concep. Bay S 1**

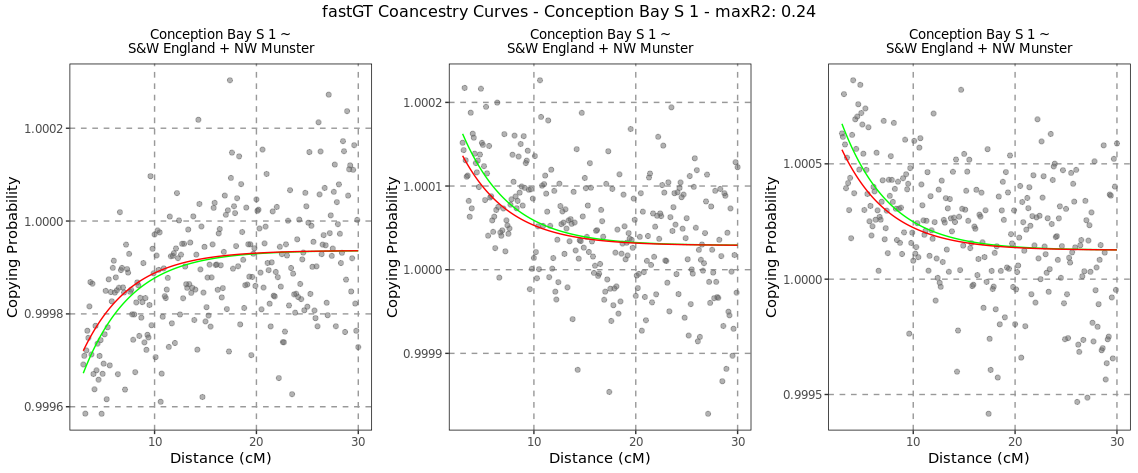

**Figure S5.14 – Bonavista Bay N 1**

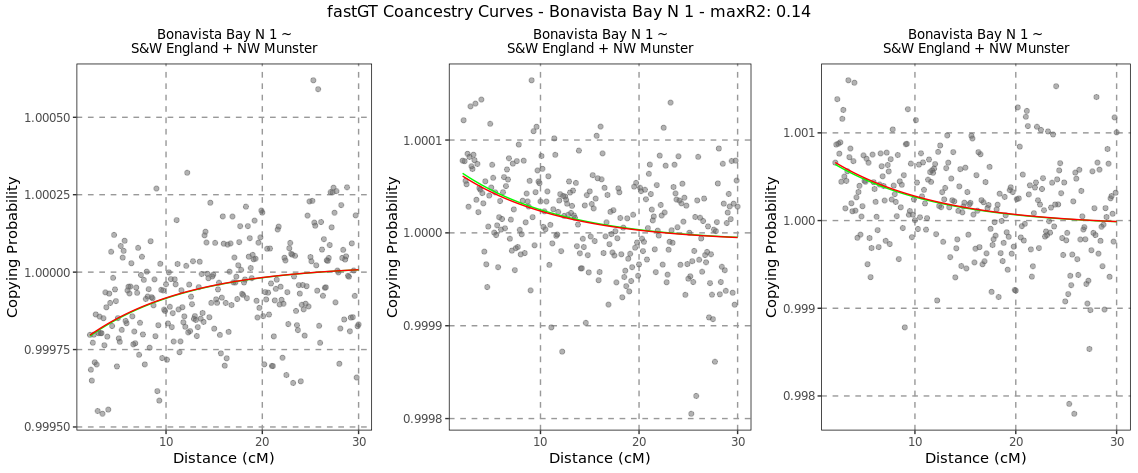

**Figure S5.15 – Bonavista Bay N 2**

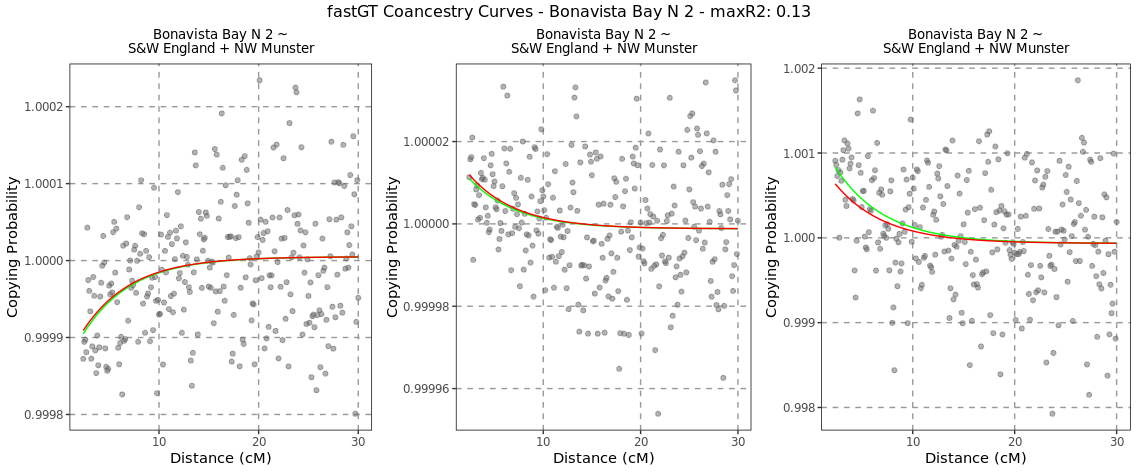

**Figure S5.16 – Bonavista Bay S**

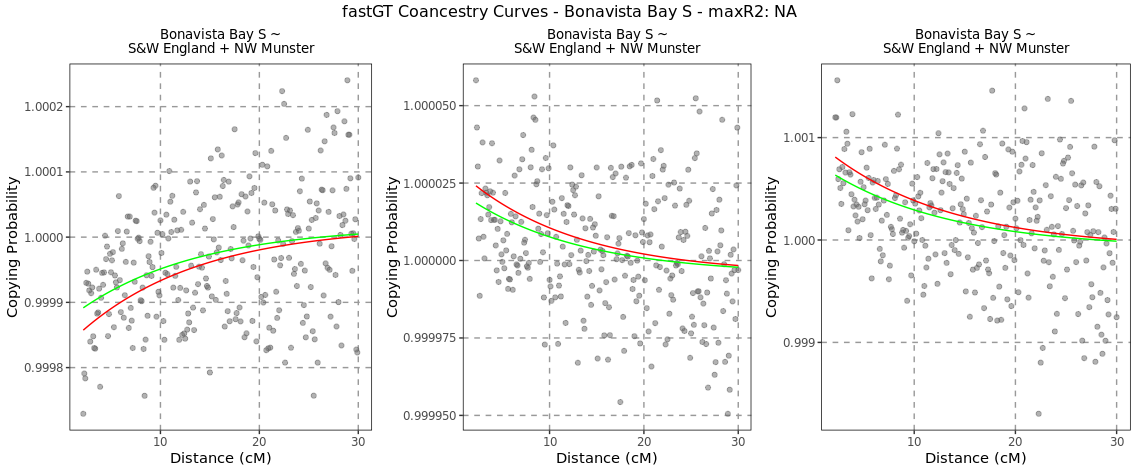

**Figure S5.17 – Conception Bay S 2**

**Figure S5.18 – Conception Bay W**

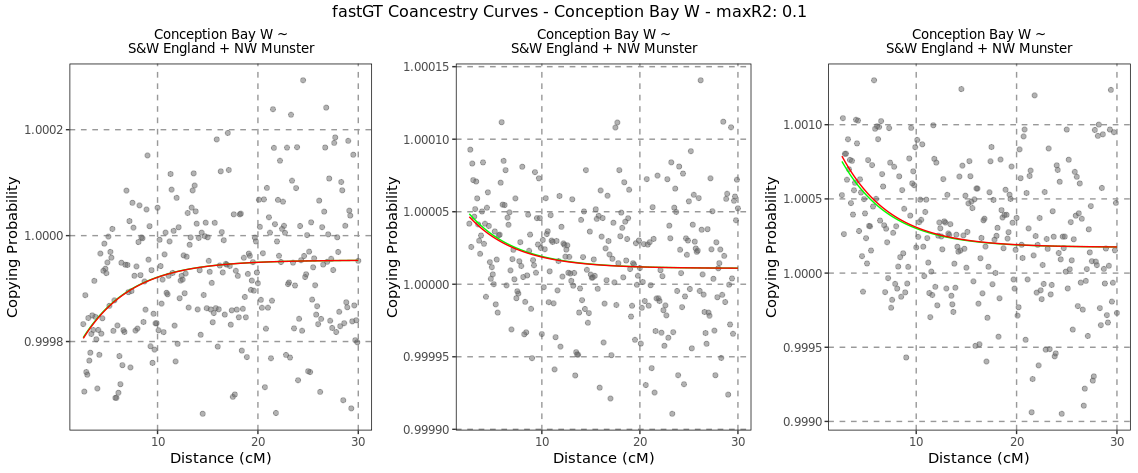

**Figure S5.19 – Trinity Bay SW**

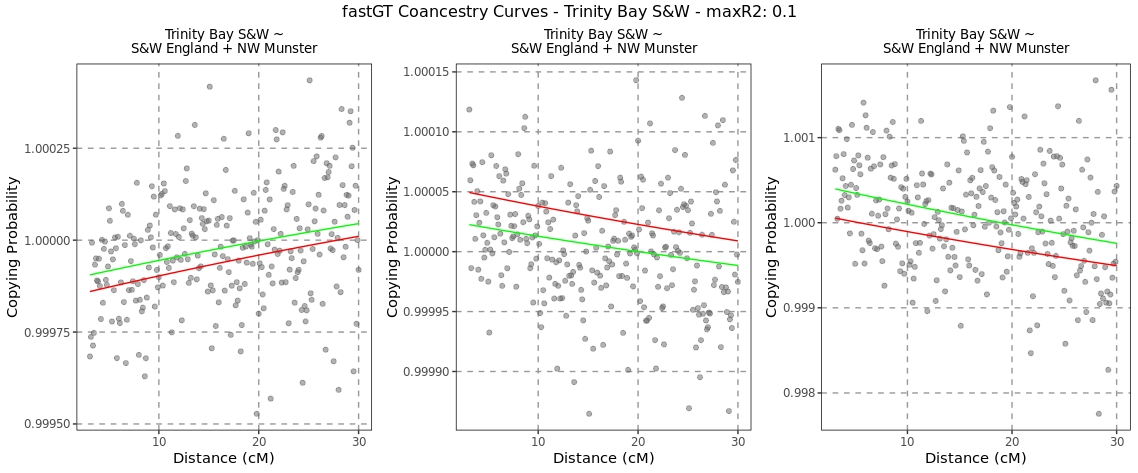

**Figure S5.20 – Trinity Bay SE**

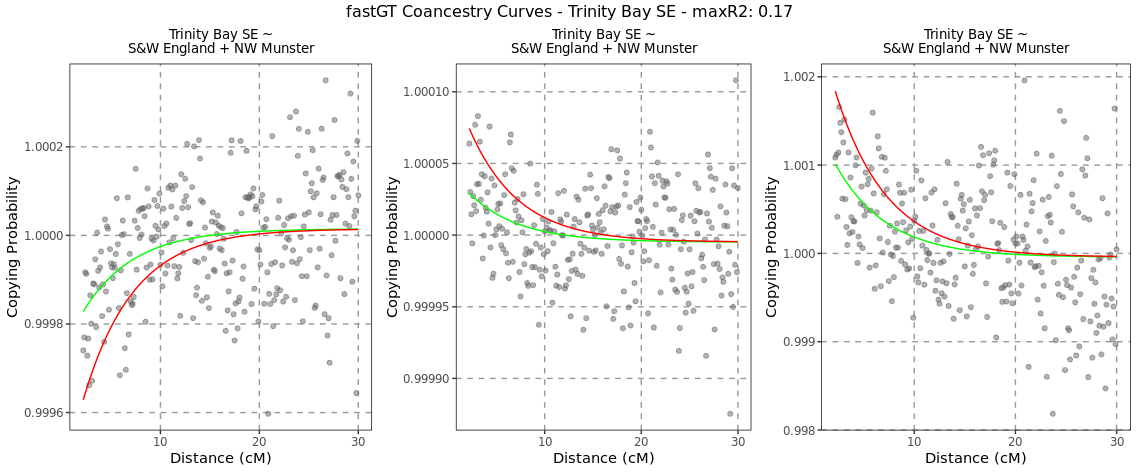

### **References**

1.	Genomes Project, C. *et al.* An integrated map of genetic variation from 1,092 human genomes. *Nature* **491**, 56-65 (2012).

2.	Li, J.Z. *et al.* Worldwide human relationships inferred from genome-wide patterns of variation. *Science* **319**, 1100-4 (2008).

3.	Alexander, D.H., Novembre, J. & Lange, K. Fast model-based estimation of ancestry in unrelated individuals. *Genome Res* **19**, 1655-64 (2009).

4.	Genomes Project, C. *et al.* A global reference for human genetic variation. *Nature* **526**, 68-74 (2015).

5.	Lawson, D.J., Hellenthal, G., Myers, S. & Falush, D. Inference of population structure using dense haplotype data. *PLoS Genet* **8**, e1002453 (2012).

6.	McInnes, L. & Healy, J. UMAP: Uniform Manifold Approximation and Projection for Dimension Reduction. *ArXiv e-prints* (2018).

7.	van der Maaten, L.J.P. & Hinton, G.E. Visualizing high-dimensional data using t-SNE. *J. Mach. Learn. Res* **9**, 2579–605 (2008).

8.	Leslie, S. *et al.* The fine-scale genetic structure of the British population. *Nature* **519**, 309-314 (2015).

9.	Gilbert, E. *et al.* The Irish DNA Atlas: Revealing Fine-Scale Population Structure and History within Ireland. *Sci Rep* **7**, 17199 (2017).

10.	Byrne, R.P. *et al.* Insular Celtic population structure and genomic footprints of migration. *PLoS Genet* **14**, e1007152 (2018).

11.	Hellenthal, G. *et al.* A genetic atlas of human admixture history. *Science* **343**, 747-751 (2014).

12.	Team., R.C. R: A language and environment for statistical computing. *R Foundation for Statistical Computing.* (2017).
